# Stress vesicles are induced by acute mechanical force and precede the commitment of epidermal stem cells to terminal differentiation

**DOI:** 10.1101/2022.09.29.510204

**Authors:** Sixia Huang, Paola Kuri, Jonathan Zou, Adriana Blanco, Maxwell Marshall, Gabriella Rice, Stephen Prouty, Tzvete Dentchev, Miriam Doepner, Joel D. Boerckel, Brian C. Capell, Todd W. Ridky, Panteleimon Rompolas

**Affiliations:** Department of Dermatology, Institute for Regenerative Medicine, University of Pennsylvania Perelman School of Medicine, Philadelphia, PA, USA; Department of Engineering, Penn State University, PA, USA; Department of Orthopaedic Surgery, University of Pennsylvania Perelman School of Medicine, Philadelphia, PA, USA; Department of Bioengineering, University of Pennsylvania, Philadelphia, PA, USA; Institute for Regenerative Medicine, University of Pennsylvania, Philadelphia, PA, USA

## Abstract

The skin has a pronounced ability to adapt to physical changes in the environment by exhibiting plasticity at the cellular level. Transient mechanical deformations applied to the skin are accommodated without permanent changes to tissue structure. However, sustained physical stress induces long-lasting alterations in the skin, which are mediated by shifts in the fates of epidermal stem cells. To investigate this phenomenon, we implemented two-photon intravital imaging to capture the responses of epidermal cells when an acute mechanical force is applied to the live skin. We show that mechanical stress induces the formation of intracellular vesicles in epidermal stem cells, which are filled with extracellular fluid and gradually enlarge, causing the deformation of the cell nucleus. By lineage tracing analysis we demonstrate that the degree of nuclear deformation is linked to cell fate. Utilizing a fluorescent *in vivo* reporter, to capture intracellular calcium dynamics, we show that mechanical force induces a sustained increase in intracellular calcium within basal epidermal stem cells. Conditional deletion of Piezo1, a mechanosensitive ion channel, alters intracellular calcium dynamics and increases the number of stress vesicles in epidermal stem cells. Using a human skin xenograft model, we show that stress vesicles are a conserved phenomenon in mammalian skin. This study uncovers stress vesicles as key manifestations of the mechanism that regulates the fate of epidermal stem cells under conditions of mechanical stress, in which Piezo1 and calcium dynamics are also involved.

## Introduction

The mammalian skin acts as a barrier protecting the body from environmental assaults. To optimally serve this function, the skin has developed properties that allow the tissue to accommodate the body’s demands for flexibility and movement. Thus, the skin is able to optimally respond to various mechanical stressors by minimizing injuries and maintaining long term functionality^1^. The cellular and molecular mechanisms that allow the skin to adapt to mechanical stress while maintaining its original phenotypic properties, even after acute deformations, have not been fully resolved^2–4^. The skin consists of three major cellular layers, the epidermis, dermis and hypodermis. It is well established that the mechanical properties of the skin are primarily dictated by the dermis, which hosts various cell types interspersed within a rich extracellular matrix, consisting of collagen and elastin fibers^5–10^. However, relatively little is known for how keratinocytes - the building blocks of the skin epithelium - internalize mechanical cues to regulate their activity and support tissue homeostasis.

Previous studies have uncovered mechanisms that describe how individual cells perceive external forces and translate them to signaling cascades, propagated from the cell surface to the nucleus and the genome^11–16^. This fundamental physiological process enables cells to induce a range of biological effects, including changes in cell fate^17–19^. Mechanotransduction at plasma membrane and the role of the nucleus as a mechanosensitive organelle are now well documented^20–28^. However, the sequence of sub-cellular events that link the plasma membrane, with the cytoskeleton and the nucleus when acute mechanical force is exerted on a live tissue, such as the skin epidermis, are less understood. The epidermis exhibits one of the highest cellular turnovers in the mammalian body, fueled by the activity of keratinocytes in the basal layer^29–32^. These cells form adhesions with the basement membrane as well as with their terminally differentiated progeny that makes up the suprabasal layers of the epidermis. Since basal keratinocytes have the exclusive ability to proliferate or commit to terminal differentiation, changes in the environment that can influence their fate may have a profound effect on tissue homeostasis. A major aim of this study is to resolve how epidermal cells respond to mechanical stress *in vivo* and how these responses may regulate their fate.

Changes in intracellular calcium dynamics are recognized as one of the earliest events in the mechanotransduction process, which can initiate the activation of downstream signaling pathways^33–38^. Recent studies have made substantial progress on visualizing the mechanosensitive calcium responses in different tissues *in vivo* and *in vitro*^26,36,39–44^. The calcium dynamics in epidermal keratinocytes have not been resolved under conditions of mechanical stress *in vivo*. Piezo1 is an ion channel and a bona fide mechano-sensor ^39,41,45–49^. Piezo1 is involved in vascular development and physiology ^40,50,51^, cell volume maintenance in erythrocytes^52–54^, and cell differentiation and proliferation^24,55–57^, among other functions. Physical tension across the cell membrane is sufficient to activate Piezo1, which allows for flux of calcium across membranes ^24,58,59^. Resolving the role of Piezo1 in the mechanobiology of epidermal keratinocytes is important given the well-established role of calcium in keratinocyte differentiation.

Here, we employed intravital imaging of mouse skin to capture in real time the cellular events induced by physical force applied directly to the live tissue. We discovered previously unrecognized subcellular processes in basal layer epidermal keratinocytes as a direct result of acute mechanical stress. Using single cell lineage tracing, we provide evidence that link these events to changes in cell fate. Furthermore, our data support a role for Piezo 1 as a critical component of this molecular mechanism that translates physical cues to cellular behaviors via calcium signaling.

## Results

### Mechanical stress induces intracellular changes in basal epidermal cells

To test how keratinocytes respond to physical stressors at the single cell lever we designed three independent assays to exert acute mechanical force onto the mouse skin epidermis. These included applying negative (suction) or positive (compression) pressure *in vivo*, as well as lateral tension (stretching) ex vivo. These assays were designed to integrate with our two-photon intravital imaging platform to directly visualize the morphological and structural changes that mechanical stress induces to skin keratinocytes within their native environment (Fig. S1A). We combined these with select mouse reporter lines to visualize different cellular structures, including the membrane (*R26-mTmG)* and nucleus (*K14-H2BGFP)* (Fig. S1B). By performing live imaging, during or immediately after the application of an acute mechanical force to the skin, we captured a formation, and rapid growth, of large vesicular structures in the cytoplasm of keratinocytes (Fig. 1A-C). This phenomenon was most prevalent in the stem cell population that resides in the basal layer of the epidermis.

**Figure 1.**
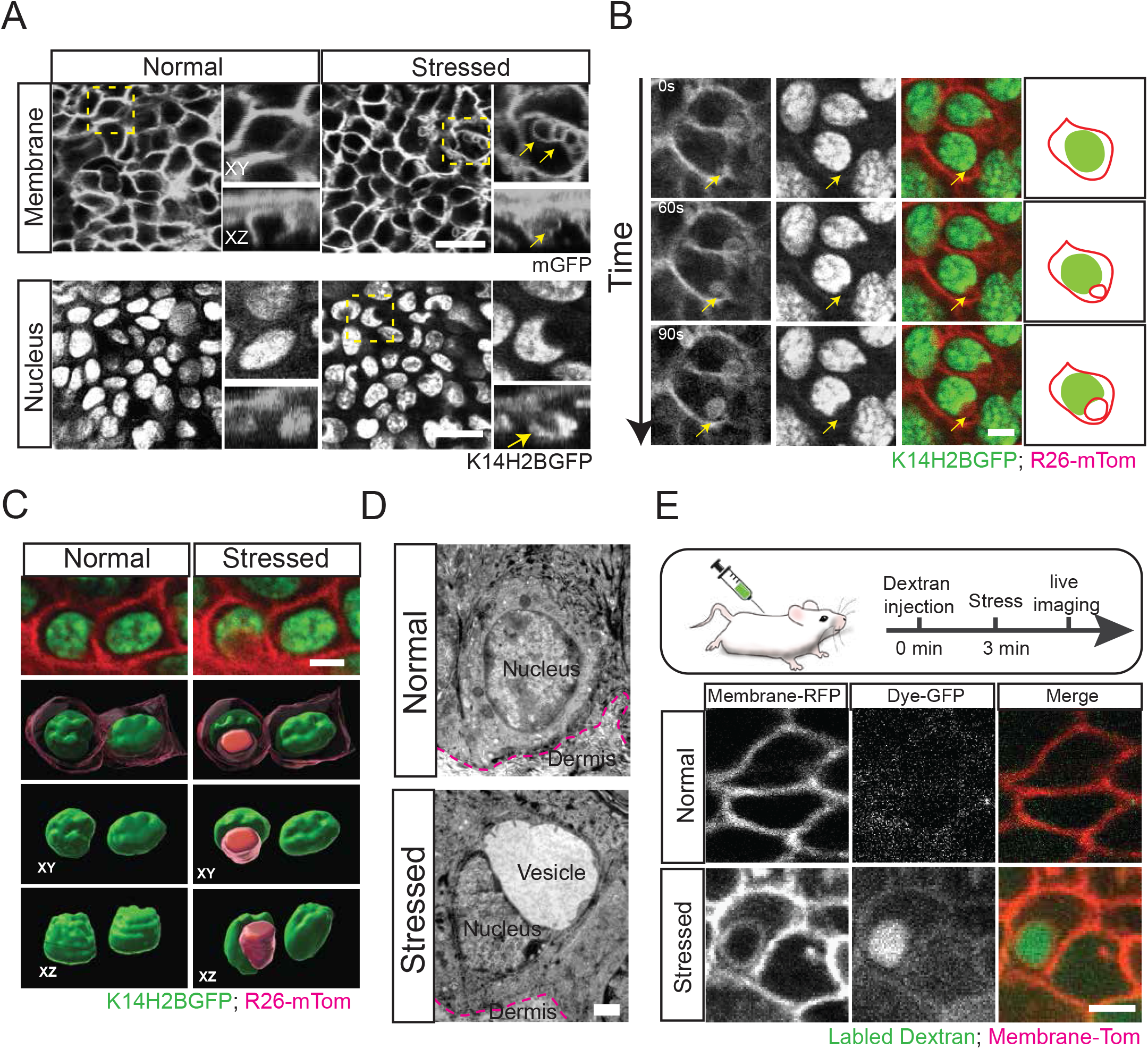
Live imaging of epidermal cell behaviors in response to mechanical force. (A) Representative images of stress vesicles (yellow arrows) in adult mouse skin after application of negative pressure, captured by two-photon intravital imaging. Insets show magnified XY and XZ plane views of outlined yellow-dotted boxes. Scale bars, 10 µm. (B) Representative time frames of stress vesicle growth in mouse skin during application of positive pressure. Scale bars, 2 µm. Also see <u>Movie S1</u>,<u>Movie S2</u>. (C) 3-dimensional renderings of the same basal epidermal cells before and after mechanical force application. The cell nucleus is represented in green, plasma membrane in magenta, and stress vesicle in red. Scale bars, 2 µm. (D) High-resolution electron micrograph of mouse basal epidermal cells. A large vesicle can be seen in the cytoplasm of the stressed cell. A red dashed line demarcates the border between the epidermis and dermis. Scale bars, 1 µm. (E) Experimental strategy to label extracellular fluid before mechanical force application and a representative example of a stress vesicle filled with Dextran-Alexa488. Scale bars, 2 µm.

Time-resolved live imaging using dual reporter mice to concurrently visualize the cell membrane and nucleus of basal layer keratinocytes captured a dynamic process, which involved the inward budding of the plasma membrane into vesicles and their gradual growth, eventually leading to the deformation of the cell nucleus (Fig. 1B, C). Ultrastructural analysis confirmed the vesicular nature of the stress-induced intracellular structures (thereafter referred to as “stress vesicles”), and showed no obvious damages in the plasma membrane, suggesting that the cells which exhibit these remain structurally intact (Fig. 1D).To test whether the formation of the stress vesicles is driven by external hydrostatic pressure during the application of an acute mechanical force, we injected the skin hypodermically with the water soluble and biologically inert fluorescent labeled dextran, to trace the movement of extracellular fluid (Fig. 1E). By live imaging, we observed that labeled dextran was concentrating in the newly formed stress vesicles confirming our hypothesis (Fig. 1E).

To further characterize the molecular composition of the stress vesicles, we performed histological analysis using markers for components of the membrane, cytoskeleton, and cell adhesion apparatuses (Fig. 2A-E). Our data show that stress vesicles are enclosed in a lipid bilayer that can be labeled by wheat germ agglutinin (WGA; Fig. 2B). A high concentration of Myosin, Vinculin and F-actin indicated the involvement of the cytoskeletal machinery in the induction or growth of the stress vesicles (Fig. 2C-E). Cell membrane dynamics is associated with the organization of contractile actin and myosin-rich stress fibers. To investigate whether actomyosin activity is required for the formation stress vesicles, we performed the stress-inducing *in vivo* assays using inhibitors against actin polymerization (Cytochalasin)^60^, and non-muscle myosin (Blebbistatin)^61^, and documented their effect in basal keratinocytes by live imaging (Fig. 2F). Under these conditions, Cytochalasin did not have a measurable effect on the formation of stress vesicles. However, injection of Blebbistatin one hour before the application of acute mechanical force reduced the number of stress vesicles by ~ 50% (Fig. 2G). Taken together, our data implicate the contractile actomyosin cytoskeleton in modulating the formation and/or growth of stress vesicles in epidermal stem cells (Fig. 2H).

**Figure 2.**
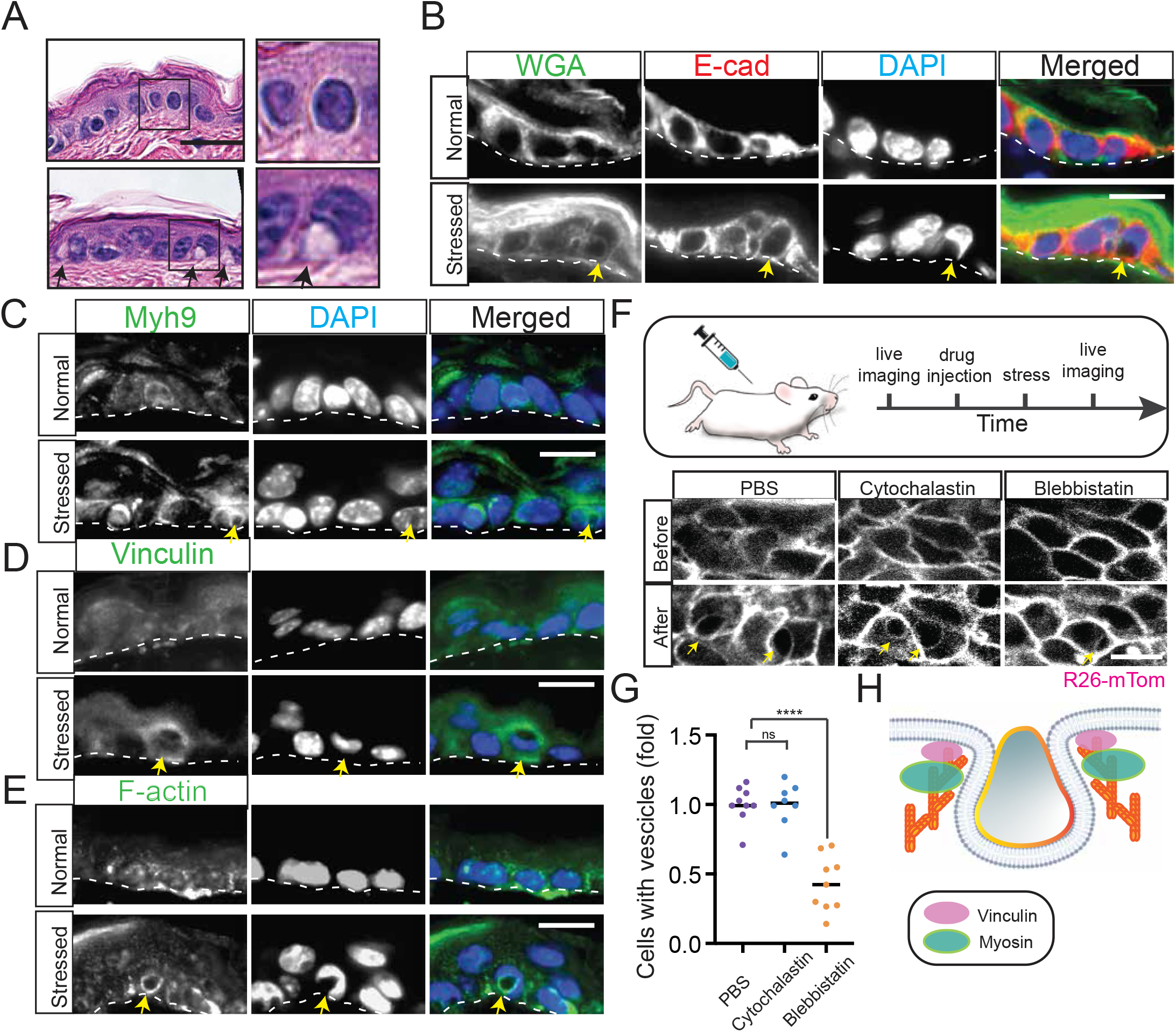
Cytoskeletal dynamics in the formation of stress vesicles. (A) H&E staining of normal and stressed skin samples. Scale bars, 10 µm. (B) Immunofluorescence staining of WGA (green), E-cadherin (red) in normal and stressed epidermis. Scale bars, 5 µm. (C), (D), and (E) Immunofluorescence staining of Myh9, vinculin, and F-actin. Scale bars, 5 µm. (F) Strategy to visualize the effect of pharmacologic inhibition of actomyosin contractility to stress vesicle formation. Representative images of the same epidermal cells before and after treatment. Yellow arrow indicates cells in the basal layer with stress vesicles. Scale bars, 2 µm. (G) Quantification of the relative ratio of stress vesicle formation in epidermal cells. n = 9 (PBS), n = 8 (Cytochalasin), n = 9 (Blebbistatin), p = 0.9116 (ns) and p < 0.0001 (****). (H) Graphical representation of actomyosin activity during stress vesicle formation.

### Stress vesicles are associated with changes in stem cell fate

Epidermal homeostasis is maintained by balanced proliferation and differentiation of basal layer keratinocytes. To investigate whether the appearance of stress vesicles in epidermal stem cells is associated with cell fate decisions, we implemented a lineage tracing approach by longitudinal live imaging. To selectively mark the stem cells in the interfollicular epidermis we used a photo-activatable *in vivo* reporter (*K14H2BPAGFP)* that generates a strong nuclear signal upon activation and which retains the long half-life of the Histone H2B moiety (Fig. 3A). First, we labeled a small group of cells, located in the basal layer of the interfollicular epidermis. We then applied a mechanical force to the skin and re-imaged the same group of cells immediately after as well as 24 hours later (Fig. 3B). We reasoned that by quantifying changes in the number and location of the labeled cells we could infer their tendency to proliferate (increase in cell number) and/or terminally differentiate (delamination from the basal layer). In the treated skin we found that the labeled group had increased in size, indicating higher proliferation compared to skin that did not experience stress (Fig. 3C). Additionally, more cells appeared in the suprabasal layers of the epidermis, 24 hours after treatment, suggesting an increase in delamination and cell differentiation after mechanical stress (Fig. 3D).

**Figure 3.**
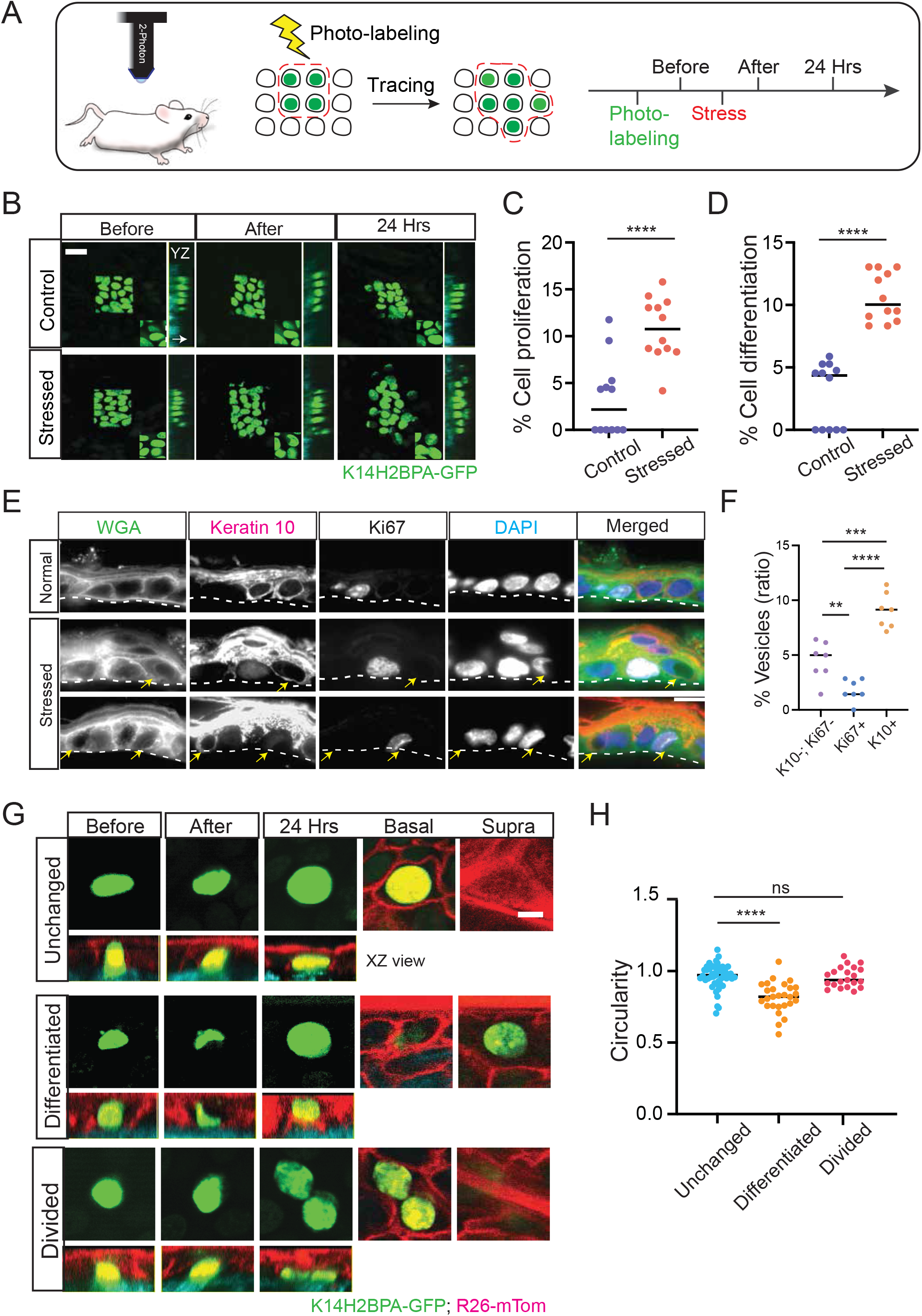
Nuclear deformation links stress vesicles to cell differentiation. (A) Genetic alleles and strategy to lineage trace epidermal cells by *in vivo* photo-labeling. Prior to mechanical force application, equivalent groups of epidermal basal cells were scanned to activate the H2B-PAGFP reporter. The same areas were re-imaged to track changes in the population (B) Representative time frames of photo-labeling and tracking after mechanical force application. Yellow arrows show the direction of cell movement when cells differentiate from the basal layer to the suprabasal layer of the epidermis. Scale bars, 10 µm. (C and D) Quantification of the relative fraction of cell proliferation or differentiation of tracked epidermal cells 1 day after mechanical force applications. n = 248 cells (normal) and 250 cells (stressed) from 3 mice, p < 0.0001. (E) Representative immunofluorescence images of stress vesicle formation in K10+, Ki67+, and K10-/Ki67-basal epidermal cells. Yellow arrow indicates cells in the basal layer with stress vesicles. A white dashed line demarcates the border between the epidermis and dermis. Scale bars, 5 µm. (F) Quantification of the ratio of stress vesicle formation in three different groups of basal epidermal cells after mechanical force application. n= 5 mice, p = 0.0042 (**), p < 0.0001(****), and p = 0.003 (***). (G) Representative time frames from live imaging lineage tracing, capturing different degrees of nuclear deformations and the subsequent fates of the epidermal cells in which they develop. Labeled epidermal cells were categorized into three groups based on their cell fate: undifferentiated cells (remain in the basal layer), differentiated cells (delaminate and move into suprabasal layer), and proliferated cells (generate two daughter cells within the basal layer). Scale bars, 2 µm. (F) Quantification of nuclear deformation as a function of cell fate. n= 48 (undifferentiated) cells, 27 (differentiated) cells, and 21 (proliferated) cells from 3 mice, p < 0.0001, p = 0.7760, and p = 0.9612.

Furthermore, in the time-resolved live imaging experiments, we observed that stress vesicles disappeared three hours after the release of the mechanical force (Fig. S2A). To test whether proliferation is indirectly triggered by cell differentiation under conditions of acute stress, we isolated actively dividing cells (Ki67+) and analyzed the stage of their cell cycle by flow cytometry. We noticed the cell cycle of proliferating cells was shifted from G1 to the G2/M phase after treatment confirming an increased activation of the epidermal progenitors (Fig. S2A-G). Next, we tested whether the changes we observed in the cellular turnover and the fates of epidermal stem cells are due secondary effects of tissue damage and cell death induced by acute mechanical stress. Our data show no obvious structural damage to the treated epidermis, including an intact epithelium across the terminally differentiated suprabasal layers as well as an unremarkable skin dermis (Fig. S3A-C). Moreover, we did not observe intracellular vesicle formation in hair follicle epidermal cells or changes in the hair cycle after we applied a mechanical force to the skin (Fig. S3D-E). We also did not observe changes in the skin immune environment after the application of a mechanical force (Fig. S3F-G).

The live imaging data showed that basal epidermal cells do not behave in a uniform way, in response to mechanical stress. This prompted us to investigate whether the formation of stress vesicles is correlated with the differentiation status of epidermal stem cells. To test this, we categorized basal epidermal cells into three discrete groups: 1) relatively quiescent epidermal stem cells (Krt10-/Ki67-basal keratinocytes), 2) transit amplifying cells (Ki67+ basal keratinocytes), and 3) committed differentiating cells (Krt10+ basal keratinocytes). Our analysis showed that Krt10+ basal layer keratinocytes show a higher rate of stress vesicle formation, compared to the other two groups (Fig. 3E, F). Taken together, these data link stress vesicles to the early differentiation state of basal epidermal cells.

### Stress-induced changes in nuclear architecture precede stem cell differentiation

Our data suggest that acute mechanical stress induces an immediate increase in the cellular turnover of the epidermis, which is likely driven by the activity of the stem cells in the basal layer. To test this hypothesis, we analyzed a possible link between the size of the stress vesicles - and the degree of nuclear deformation induced by their growth - to a particular cell fate. For this, we photo-labeled individual basal epidermal cells and traced their activity after applying mechanical force to the skin (Fig. 3G). By analyzing the geometric characteristics of the nucleus, we found that cells with a highly deformed nucleus exhibited a distinct tendency to differentiate and leave the basal layer. In contrast, cells without dynamical intracellular changes, such as no stress vesicles and intact nuclei, either divided or remained unchanged within the basal layer (Fig. 3H). Previous studies have shown that cells in the basal layer of the epidermis experience major changes in cell shape, cell adhesiveness and cytoskeletal architecture during the different phases of cell the cycle, and that these cellular changes are also associated with the responses of cells to external mechanical stimuli^62–65^.

Interestingly, the pro-differentiation phenotype of epidermal stem cells as result of mechanical stress, appears to be linked to the degree of the nuclear deformation rather than to changes in nuclear volume (Fig. S4A-E). Previous studies demonstrated that the cell exhibits a certain level of elasticity when mechanical force is applied^15,26,28,66–69^. The nuclear lamina is critical for the structural stability of the nucleus and plays a key role in chromatin organization and genome integrity. Lamin A/C is a key component of the nuclear lamina and evidence suggests that cells with higher nuclear compression express low levels of Lamin A/C and vice versa ^70–72^. After applying mechanical stress, in epidermal cells with deformed nuclei, the shape of nuclear lamina (Lamin A/C staining) matched that of the nuclear content (H2BGFP signal). However, we did not observe a change in the overall levels of Lamin A/C in cells with stressed vesicles, compared to control or to cells with intact nuclei (Fig. S4F). These data suggest that stress vesicles and their impact to cell fate likely work independently to the stiffness of the nuclear envelope.

### Epidermal cells sense stress through persistent intracellular Ca2+

Calcium is a key mediator of cellular mechanotransduction in various systems^35,38,39,44^. Moreover, calcium is a major regulator of keratinocyte differentiation *in vivo* and *in vitro*^37,64,73^. Our observation of stress vesicles in epidermal stem cells and their link to cell differentiation indicated that calcium may be implicated in this pathway. To test this, we generated mice in which epidermal cells express the calcium reporter GCaMP6s. Thus, we aimed to capture the calcium dynamics in the skin, under physiological and stressed conditions, by live imaging (Fig. 4A). In the adult epidermis, keratinocytes are organized in layers which reflects their state of differentiation. This functional segregation of stem cells and their committed progeny is mirrored by a calcium gradient from the basal to the terminally differentiated suprabasal layers (Fig. S5B)^74,75^. In untreated skin, live imaging of the mouse ear skin captured complex patters of intracellular calcium fluxes, exhibited specifically by basal layer keratinocytes, consistent with previous observations (Fig. 4B)^64^.

**Figure 4.**
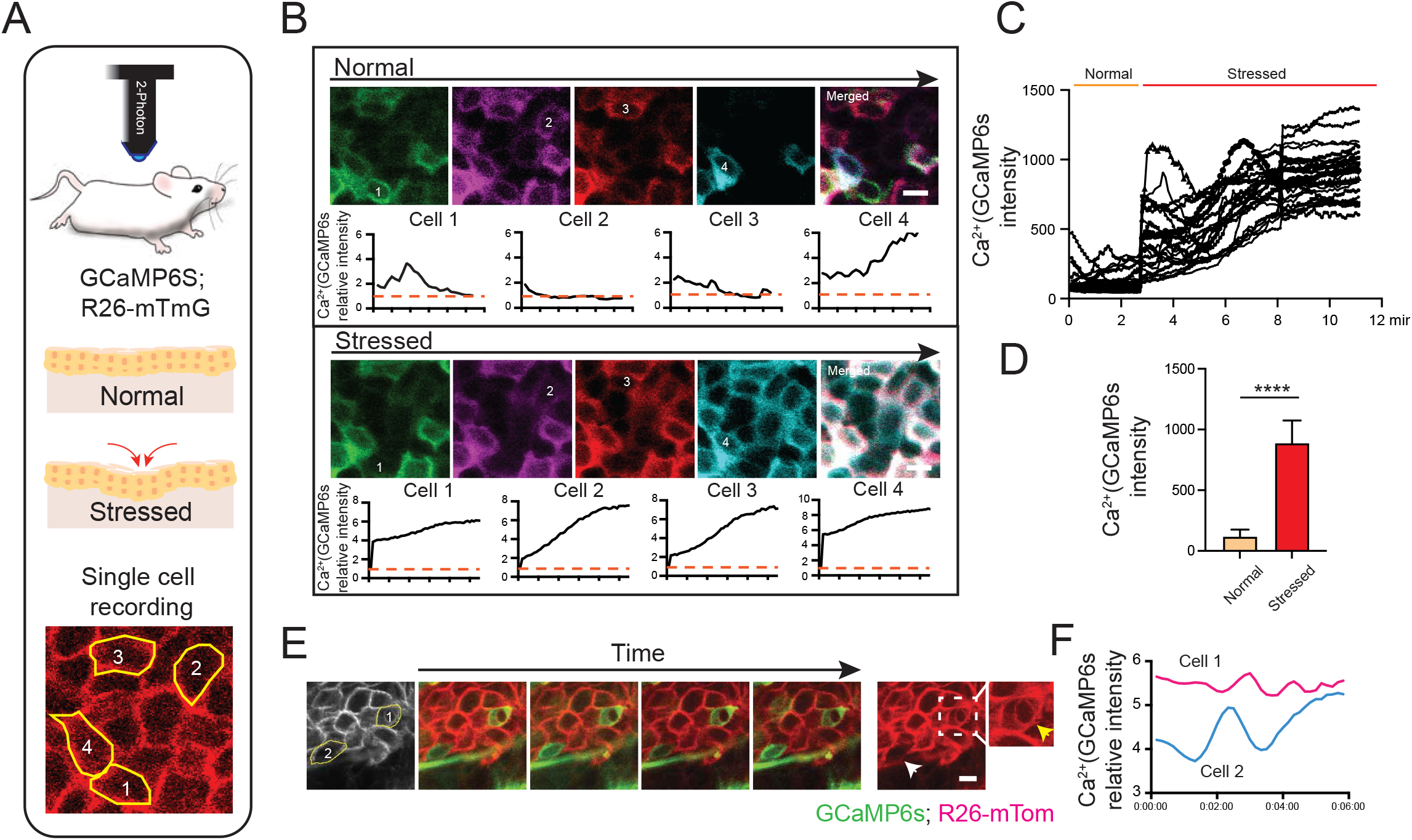
Mechanical stress alters intracellular calcium dynamics. (A) Schematic of calcium imaging of individual epidermal cells while applying compression on the live mouse skin. (B) Five representative time frames of *in vivo* calcium imaging of basal epidermal cells before (top panels) and after (bottom panels) mechanical stress. Each frame was pseudo-colored and projected into the last frame (right). Note the initial intracellular calcium fluxes in four highlighted epidermal cells and the persistence of cytoplasmic calcium in the same cells after applying compression on the skin. Scale bars, 2 µm. Graphs show relative intensity of calcium signal as a function of time for each highlighted cell. (C) Quantification of calcium signals at rest and under stress. Each line presents one cell, n=24 cells from 1 mouse, this experiment was repeated in 3 different mice. (D) The cumulative sum of calcium signals at baseline and under stress. n = 24 cells, p < 0.0001. (E) Representative examples of calcium dynamics in the cells with or without stress vesicles Scale bars, 2 µm. (F) Quantification of the relative calcium intensity as a function of time for two representative cells. In this example only Cell 1 formed a stress vesicle.

To detect how epidermal stem cells modulate intracellular calcium in response to an acute mechanical stimulus, we applied a compressive force to the mouse ear skin and directly visualized the calcium dynamics in the affected epidermal cells. Live imaging data revealed a tissue-wide response, with persistent high calcium signal in the cytoplasm of basal layer keratinocytes, shortly after the application of a mechanical force (Fig. 4B-D). Applying the same level of compression for a longer time, lead to a gradual decrease in intracellular calcium, indicating a level of cellular adaptation to the new mechanical environment. Importantly, we observed that stress vesicles preferentially formed in cells that exhibited persistent high cytoplasmic calcium as opposed to those in which cytoplasmic calcium fluxed (Fig. 4 E, F). Taken together, the data provide a link between acute mechanical stress, intracellular calcium and stress vesicles in epidermal stem cells.

### Piezo1 deletion increases stress vesicle formation

Piezo1 is a mechanosensitive ion channel which enables cells in various tissues to perceive and respond to mechanical stimuli^54^. Even though Piezo1 is a general cation channel, it influences cell activity primarily by controlling calcium flows across membranes. Based on our data, we hypothesized that Piezo1 may be involved in the same pathway that modulates stress vesicles and their effect on epidermal stem cell fate. To test this *in vivo* we conditionally ablated Piezo1 gene expression in adult mice using a strong basal driver (p63CreER; Fig. 5A). Piezo1 is expressed in basal layer keratinocytes and is effectively eliminated in the Piezo1 conditional knockout (Piezo1 cKO; Fig. S5B). Under physiological conditions, there were no apparent differences in skin morphology between Piezo1 cKO mice compared with wildtype mice. We then investigated whether epidermal cells with Piezo1 gene deletion differ in their response to mechanical stress. Compared to wildtype mice, cells in Piezo1 cKO exhibited a marked increase in the number of stress vesicles under similar conditions of mechanical stress, suggesting that Piezo1 is involved in this process (Fig. 5B, C).

**Figure 5.**
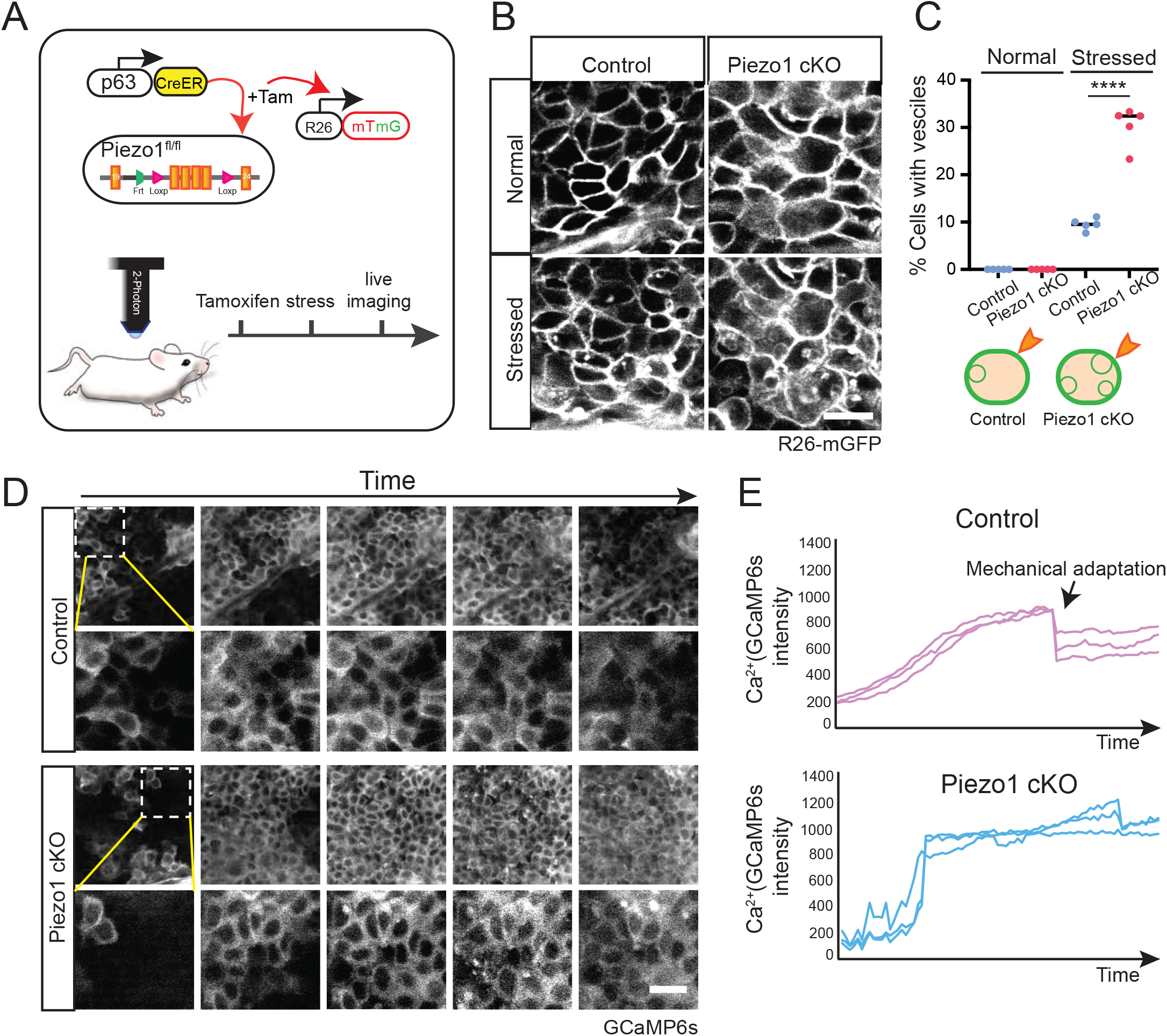
Piezo1 ablation exaggerates stress vesicle formation. (A) Schematic of experimental design to conditionally ablate expression of Piezo1 in basal epidermal cells and observe the effect on stress vesicle formation. (B) Representative images of basal epidermal cells, from control and Piezo1 conditional knockout (cKO) mouse back skin, visualized before and right after mechanical force application. Scale bars, 10 µm. (C) Quantification of the fraction of basal epidermal cells with stress vesicles. n= 5 pairs of control and Piezo1 cKO mice, p < 0.0001. (D) Representative images of calcium signals at the plane of basal layer of the skin, from control and Piezo1 cKO mice, taken immediately after mechanical force application. Scale bars, 20 µm. (E) Quantification of calcium dynamics in 3 representative basal epidermal cells from Control and Piezo1 cKO mice under stressed conditions.

An acute mechanical force exerted on the tissue is predicted to propagate across the cellular membranes on which Piezo1 is localized, leading to channel activation and flow of calcium into the cytoplasm from extracellular and/or intracellular stores. To investigate this in the skin epidermis, we bred mice that allowed us to image calcium dynamics in the presence or absence of Piezo1. Under baseline conditions, we observed no marked differences in the calcium fluxes that we previously documented in the basal layer of the epidermis (Fig. S5C, D). Furthermore, both control and Piezo1 basal keratinocytes exhibited an increase in intracellular calcium when an acute mechanical force was applied to the skin (Fig. 5D). However, the kinetics of the increase in intracellular calcium under the same mechanical stress were markedly different between the two groups. Specifically, while in wild type keratinocytes the intracellular calcium increased gradually, in Piezo1 cKO cells the change was abrupt (Fig. 5E). Interestingly, Piezo1 cKO cells also did not recover in the same manner as wild type cells and experienced persistent high intracellular calcium, long after the removal of the stress stimulus (Fig. 4E, 5D and 5E). These results demonstrate that basal epidermal cells respond to mechanical force by increasing intracellular calcium and that the kinetics of calcium influx partially dependent on Piezo1activity.

To directly test whether Piezo1 channel activation could prevent intracellular vesicle formation, we intradermally injected the Piezo1-selective activator Yoda1 before application of a mechanical force to the mouse skin. As predicted, Yoda1 induced a robust increase in calcium influx (Fig. S5E). More importantly, Yoda1 treatment suppressed the formation of stress vesicles in live skin under conditions of mechanical stress (Fig. S5F, G). Conversely, we found that treatment with a calcium channel inhibitor (GsMTx4) did not prevent stress vesicles from forming (Fig. S5F, G), which further confirmed that Piezo1 is a key mediator of stress vesicles.

To further dissect the molecular pathways involved in epidermal stem cell stress response, we performed RNA sequencing analysis on basal layer keratinocytes isolated from the interfollicular epidermis of wildtype and Piezo1 cKO mice. Gene expression analysis highlighted genes involved in barrier function and cell adhesion, which where downregulated in the Piezo1 cKO (Fig. S7A, B). Interestingly, we found that expression of genes implicated in immune activation were upregulated in Piezo1 cKO keratinocytes compared to control (Fig. S7B). The gene signature for development pathways primarily included TGF-beta^76^, PKC^77,78^, and calcium related genes: all signaling pathways believed to be involved in mechanotransduction and cell fate determination. To test whether Piezo1 deletion also affects epidermal integrity and barrier function, we applied a persistent mechanical force to the live mouse skin. After 10 min of applied negative pressure, Piezo1 cKO skin exhibited extensive blistering compared to control mouse back skin (Fig. S5B, C). Taken together these data indicate that that the mechanosensitive ion channel Piezo1 plays a protective role in maintaining skin integrity and preventing tissue rupture from acute mechanical stress.

### Stress vesicles are a conserved phenomenon in mammalian skin

Despite the molecular and functional similarities, the structural organization and mechanical properties of mouse skin are markedly different than human. We wondered whether the phenomena we observed in the mouse skin are also relevant to human skin physiology and if human keratinocytes exhibit stress vesicles in response to mechanical stress. Conventional cell culture assays cannot sufficiently recapitulate the biomechanical environment of the intact skin. To address this challenge, we devised a strategy that would allow us to replicate the mechanical stress conditions in a human xenograft model and use live imaging to directly visualize the responses of human keratinocytes after application of an acute mechanical force (Fig. 6A). We incorporated fluorescent reporter genes in the human primary keratinocytes that we used to establish the organotypic culture, in order to illuminate the nucleus, cell cortex and cell membrane. After the engineered human skin was fully stratified, we grafted it in mice and tested for long-term stable integration and viability (Fig. 6B). Live imaging of the human skin xenograft validated the presence of labeled keratinocytes in the tissue, capturing the columns of differentiating cells (Fig. 6D). Histological analysis showed that the engineered xenograft displayed the distinct morphology and cellular organization of human skin, including the presence of ridge-like structures and the vastly higher number of differentiated layers compared to the surrounding native mouse skin (Fig. 6D). We then applied the same mechanical stressors to the human skin xenograft and imaged their effect to keratinocytes by live imaging. Stress vesicles with similar appearance and abundance formed consistently in the human skin xenograft confirming that this is a conserved phenomenon. Taken together our data illuminate a previously unrecognized mechanism that mammalian skin has developed to respond and adapt to acute mechanical stress (Fig. 6E).

**Figure 6.**
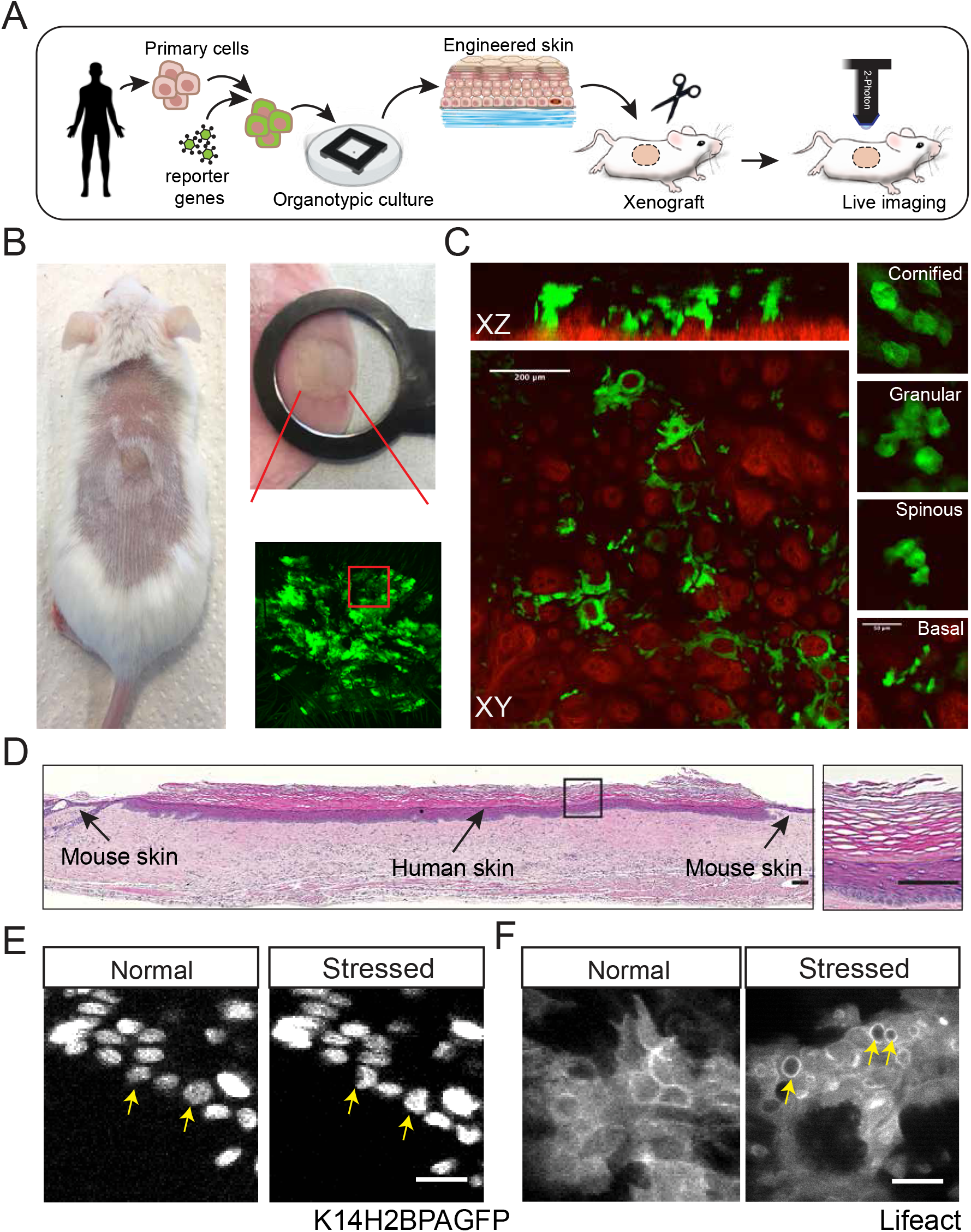
Formation of stress vesicles in a human skin xenograft. (A) Schematic of the experimental design to generate a human skin xenograft for intravital imaging. (B) Representative examples of a stable human skin xenograft in the mouse. Magnified views show the mounting (top) and the wide field epifluorescence image of the xenograft, acquired by live imaging. (C) Representative low and high magnification serial optical sections and corresponding orthogonal views of the human skin xenograft, by two-photon microscopy. Note the ridge-like structures in the epidermal-dermal junction (Second Harmonic Generation in red), the epithelial clonal columns (K14-GFP) and the distinct multilayered organization recapitulating the human skin epidermis. Scale bars, 200 µm and 20 µm. (D) H&E stained histological section of the entire length of the human skin xenograft. Scale bars, 100 µm. (E-F) Representative examples of stress vesicles forming in basal epidermal cells of the human skin xenograft. Note the deformation of the nucleus that recapitulates what we observed in the mouse skin. Scale bars, 10 µm.

## Discussion

Tasked with maintaining a barrier that shields the body from the outside environment, the skin is an organ uniquely exposed to physical stressors. To maintain structural integrity and deal with cellular attrition under these harsh conditions, the skin epidermis displays a constant cellular turnover that is driven by the activity of stem cells in the basal layer of the stratified tissue. While the skin can appear phenotypically normal after experiencing an acute mechanical force, the underlying cellular mechanisms that enable this robustness are not fully understood. Leveraging intravital imaging to directly observe the responses of keratinocytes within their native tissue environment at the single cell level, we captured a conserved sequence of intracellular events that culminated in the formation of stress vesicles, in response to mechanical stress. We provide evidence that link stress vesicles with intracellular calcium dynamics and the mechanosensitive channel Piezo1. Moreover, we demonstrate that stress vesicles induce profound changes inside the cell, leading to the deformation of the nucleus. More importantly, we show that the degree of nuclear deformation can predict whether a cell will commit to terminal differentiation, directly implicating this phenomenon to the regulation of cellular fate in the adult epidermis. The physiological significance of these processes is not immediately apparent; however, the data may point to several plausible hypotheses.

The adult skin is made up of three main layers: the epidermis, dermis, and the subcutaneous layer. It is also perfused with extracellular fluid that acts as a bridge between cells^79,80^. Under normal conditions the hydrostatic pressure of extracellular fluid is equilibrated across the tissue. An acute mechanical force applied to the skin can disrupt this balance leading to rapid local increases in hydrostatic pressure. These can lead to the displacement or shearing of cell adhesion molecules allowing high pressure fluid to enter the space between cells pushing the plasma membranes inward. There is precedence of phenomena described in vitro, as well as during development *in vivo*, that pose striking similarities to our observed stress vesicles, which are induced by fluid pressure and disruption of cell adhesion^81–85^. Our data showing that stress vesicles extracellular fluid support this hypothesis. While localized increases of hydrostatic pressure due to mechanical stress are likely triggering the invagination of the plasma membrane, our results suggest that keratinocytes may have developed mechanisms to control this process by engaging their cytoskeletal machinery, in order to minimize damage to the cell.

Epithelial cells have developed ways to translate mechanical cues into biochemical signals. Key components of the mechanotransduction machinery are adhesion complexes embedded into the plasma membrane that mediate the transmission of physical forces from the external environment to the intracellular space and the nucleus to regulate gene expression^86–89^. Mechanosensitive ion channel, such as TRP (Transient Receptor Potential) and Piezo are additional potent transducers of physical information to the cell. Their activation in response to tensile forces applied to the cell membrane allows calcium to pass across a gradient into the cytoplasm activating signaling cascades that influence cell adhesion and keratinocyte differentiation^90,91^. Piezo1, a family member of the mechanosensitive cation channels, which is also expressed in epidermal keratinocytes, has previously been implicated in mediating keratinocyte mechanical sensitivity and migration during wound healing^92,93^. Interestingly, Piezo1 activity appears to influence several intracellular processes through modulation of cytoskeletal dynamics and cell adhesion^94^. Our results show that in the epidermis conditional Piezo1 deletion deregulates intracellular calcium flux, alters the expression of genes involved in cell adhesion and membrane dynamics and amplifies the appearance of stress vesicles under acute mechanical stimulation. In this context, Piezo1 may act as a sentinel mechanism priming the cytoskeletal and adhesion apparatuses of keratinocytes to adapt in the mechanical environment of the tissue. It would be interesting to speculate an involvement of such mechanism on the regional differences observed in skin that is exposed to various degrees of mechanical stress.

The endpoint of the mechanotransduction processes is the regulation of gene expression that ultimately controls the activity and fate of cells. Several pathways have been proposed to convey physical cues to the genome to affect the function of key genes. Emerging evidence support an intriguing concept that puts the nucleus at the center of the mechanosensory apparatus. Based on this, mechanical forces that are exerted on nucleus are translated into different modes of chromatin organization that affect transcriptional activity and cell fate^95^. Nuclear mechanotransduction can be modulated through the cytoskeleton, the nuclear lamina, and nuclear actin dynamics^26,96^. Here we show that application of an acute mechanical force to the skin induces the formation of stress vesicles in epidermal stem cells, leading to the deformation of their nucleus. A correlation between the degree of nuclear deformity to the subsequent differentiation fate of the cell suggest that stress vesicles may represent and additional mode of nuclear mechanotransduction.

During the cell cycle, epidermal keratinocytes experience profound changes in cytoskeletal architecture and cell adhesiveness. These structural changes may alter the way cells respond to acute changes in their mechanical environment. Our single cell lineage tracing data reveals that cells react to mechanical stress in diverse patterns responses depending on their differentiation status. Epidermal stem cells normally express Keratins 5/14^97–99^. However, one of the earliest indications of their commitment to terminal differentiation is a switch to expression of Keratin 1/10. The keratin intermediate filament network plays key roles in organizing the architecture of plasma membrane and nucleus^97,100–103^. Moreover, intermediate filaments have recently been proposed to regulate cell fate decision acting as mediators of cellular differentiation^104^. Our data link the transition to Keratin 10 expression with the propensity of the cell to form stress vesicles under stress conditions. It is plausible that a Keratin 1/10 intermediate filament network makes a basal cell stiffer and thus more susceptible to mechanical stress. Future work may illuminate the role of intermediate filaments as key mechanotranducers and effectors of cellular fate.

Altogether, our work demonstrates that in the skin epidermis mechanical stress induces a cascade of intracellular structural changes that begin with the inward budding and growth of stress vesicles and culminate with the deformation of the nucleus and the initiation of the terminal differentiation program. Piezo1 plays a key role in this process, linking intracellular calcium dynamics to cell adhesion and the formation of stress vesicles. Our findings suggest that these conserved phenomena may be part of a robustness mechanisms to protect the tissue from structural damage and to maintain homeostasis under conditions of mechanical stress.

## Acknowledgments

We thank George Cotsarelis, John Stanley and John Seykora for their invaluable advice that guided this study. We also thank Sara Wickström and Jean-Leon Maitre for insightful discussions. We are grateful to the University of Pennsylvania Skin Biology and Disease Research-based Center (SBDRC) for analysis of tissue sections and support with the establishment of human-engineered skin xenograft. We also acknowledge the support of the Institute for Regenerative Medicine and the entire stem cell community at Penn. S.H. was supported by an IRM postdoctoral fellowship. G.R. was supported by training grant T32GM007229 from NIH/NIGMS. P.K. was supported by the American Association for Cancer Research-John and Elizabeth Leonard Family Foundation Basic Cancer Research Fellowship. P.R. was supported by grants from NIH/NEI (R01EY030599) and from the American Cancer Society (RSG1803101DCC).

## Author contributions

S.H. and P.R. conceptualized the study, designed the experiments and wrote the manuscript. S.H., P.K., A.B., M.M., G.R. and P.R. performed the experiments. J.Z. and B.C. assisted with the RNAseq and performed the bioinformatic analysis. S.P. and T.D. performed histological analysis. P.K., M.D., and T.D. assisted with the establishment of human-engineered skin xenograft. S.P. performed histological analysis. All authors discussed results and participated in the manuscript preparation and editing. P.R. supervised the study.

## Competing interests

The authors declare no competing interests.

## Data and materials availability

Data sets and reagents presented in this study are available from the corresponding author upon request.

## Methods

### Mice

All procedures involving animal subjects were performed with the approval of the Institutional Animal Care and Use Committee (IACUC) of the University of Pennsylvania. *K14CreER, R26*^*loxp- tdTom-stop-loxp-EGFP*^ (*R26-mTmG* in text), *R26-CAG-GCaMP6s, Piezo1*^*fl/fl*^ mice were obtained from The Jackson Laboratory. *p63*^*CreER*^ mice were created by J. Xu (Baylor College of Medicine. *K14-H2B-PAGFP* mice were generated by the center for Animal Transgenesis and Germ Cell Research, at the School of Veterinary Medicine of the University of Pennsylvania. All mice that were used in this study were bred for multiple generations into a Crl:CD1(ICR) mixed background. For lineage tracing experiments and Piezo1 gene deletion experiments, Cre activation was induced with intraperitoneal injections of Tamoxifen in corn oil (0.1-2 mg per 20 g body weight). Mice were induced between 6-8 weeks old and subsequent experiments were conducted at the indicated times after induction. Experiments included equal representation of males and females. There was no apparent difference in phenotype between genders. Mice were housed in a temperature and light-controlled environment and received food and water ad libitum.

### Mechanical force assays

Mice were anesthetized with an intraperitoneal injection of Ketamine / Xylazine cocktail I PBS (0.1 mL / 20 g body weight: 87.5 mg /kg Ketamine, 12.5 mg / kg Xylazine). Mouse skin was shaved and treated with depilatory cream (Nair) prior to mechanical force application. Negative pressure: A modified cutaneous suction system with a glass vacuum chamber (10 mm in diameter) was applied to generate a negative pressure on the back skin of anesthetized mice. Briefly, a vacuum of 80 Kpa was used to generate negative pressure to the mouse back skin. For monitoring the formation of stress vesicles negative pressure was applied to mouse back skin for 3 minutes before imaging. For mouse skin fragility study, the skin was carefully monitored for blistering formation during a 10 minute application of suction force. The number of blisters was counted right after treatment. Positive pressure (Compression): The ear and back skin on each mouse was compressed by using a modified pinching method, in which a metal arm connected a cover slip compressed the mounted skin. Using this approach to generate compression on mouse skin, we can visualize the responses of epidermal cells to mechanical forces by live imaging in real time. Mechanical stretching (lateral tension): Full thickness skin (2 cm in width and 3.5 cm in length) was collected from adult mouse back and put into PBS. Each skin sample was subsequently mounted onto a custom-designed tensile stretching device with a 37°C heating element that was placed onto an imaging stage. The explant was then stretched to its original dimensions before applying tensile stress. During linear stretching the explant was continually moisturized with PBS that was preheated to 37 C°. Individual skin samples were stretched to three different strain rates (low, medium, high) from 10% to 30% strain and that are quantified by the elongated length of the skin tissue after stretching.

### *In vivo* imaging

Imaging preparation and procedures were performed with the protocol that previously described^105^. Mice were anaesthetized by intraperitoneal injection of ketamine/xylazine cocktail in PBS (0.1 mL / 20 g body weight; 87.5 mg / kg Ketamine, 12.5 mg / kg Xylazine). A surgical plane of anesthesia was verified by absence of pedal reflex responses following physical stimulation and was maintained during the imaging period with 1% vaporized isoflurane in oxygen and air delivered through a nose cone. The skin was mounted on a custom-made stage with a glass coverslip placed directly against it. To maintain body temperature, mice were placed on a heating pad throughout the experiment. Image acquisition was performed with an upright Olympus FV1200MPE microscope, equipped with a Chameleon Vision II Ti: Sapphire laser. The laser beam was focused through 10X, 20X or 25X objective lenses (Olympus UPLSAPO10×2, N.A. 0.40; UPLSAPO20X, N.A. 0.75; XLPLN25XWMP2, N.A. 1.05). Emitted fluorescence was collected by two multi-alkali and two gallium arsenide phosphide (GaAsP) non-descanned detectors (NDD). The following wavelengths were collected by each detector: NDD1 419-458 nm; NDD1 458–495 nm; GaAsP-NDD1 495–540 nm; GaAsP-NDD2 575–630 nm. GFP and Tomato reporters were excited at 930 nm and their signal was collected by GaAsP-NDD1 and GaAsPNDD2, respectively. Second harmonic generation signal was generated using 930nm excitation wavelength and detected in NDD2. Serial optical sections were acquired in 2.5-3 mm steps, starting from the surface of the epidermis and capturing the entire thickness of the epidermis and a partial section of the dermis (epidermis 30 mm, dermis 60-80 mm). Multi-day tracing experiments were performed by re-imaging the same field of view at the indicated times after the initial acquisition, with vasculature and micro-tattoos used as landmarks to identify the imaging region at low magnification and clusters of hair follicles used as landmarks higher magnification. For all time-lapse videos, the live mouse remained anesthetized for the length of the experiment and serial optical sections were capture at intervals of 10 seconds. After each imaging session, the mice were monitored and allowed to recover in a warm chamber before being returned to the housing facility.

### *In vivo* photo-labeling

The procedures were performed with the protocol that previously described^106^. The pre-activated form of the H2B-PAGFP fluorescent proteins was visualized by exciting with 850 nm wavelength and emission signal was collected in GaAsP-NDD1. Excitation with 930 nm verified that no signal is emitted by the reporters before activation. Photolabeling was achieved by scanning a defined region-of-interest (ROI) at the plane of the basal layer of the epidermis, with the laser tuned to 750nm wavelength, for 5-10 s, using 5%–10% laser power. Immediately after photo-activation, a series of optical sections, with a range that includes the entire thickness of the skin, were acquired using the same acquisition settings as for GFP. Visualizing the signal of the activated form of PAGFP only within the ROI confirmed the successful photo-labeling of basal layer keratinocytes. Following the initial image acquisition immediately after photo-labelling, the same area of skin was re-imaged at the indicated times to evaluate the responses of the individual labeled basal keratinocytes or groups of labeled basal keratinocytes to mechanical forces compared to control cells or control groups.

### Hypodermal drug injection

To label extracellular fluid, 10 kDa fluorescent dextran (Alexa Fluor 488 dextran Invitrogen D22910) was injected subcutaneous into back skin prior to mechanical forces application. Briefly, single anaesthetized mouse was placed on the heat pad, fluorescent dye at a concentration of 20mg/ml and 10 µl volume in total was injected into skin dermal area. Mechanical force was applied right after fluorescent dye injection. Images were taken in green channel immediately after mechanical force application. For the inhibitor experiments, mice were locally injected with Cytochalasin D (Abcam, ab143484, 50 µM), blebbistatin (Sigma, B0560, 2 mg/kg per mouse) 1 hour before mechanical force application, PBS was used as a vehicle control. For the activation of calcium influx experiment, 5 µM Yoda1 was injected locally in back skin prior to mechanical force application, PBS diluted DMSO was used as a vehicle control. For the inhibitor of calcium dynamic experiments, GsMTx4 (R&D, 4912, 1 µM) was locally injected in the area prior to mechanical force application, and PBS was used as a vehicle control.

### Immunostaining, TUNEL assays, and RNAscope

For section staining, dorsal skins were dissected, laid flat and directly fixed with 4% paraformaldehyde in PBS for overnight at 4 C°. Fixed tissues were paraffin embedded and sectioned. Deparaffinized slides were incubated in a 1:50 dilution of Antigen Unmasking Solution (Vector Laboratories) and heated in a boiling bath for 10 min. Sections were permeabilized with 0.2% Tween 20 in PBS and incubated in blocking solution (10% normal goat or horse serum in 0.5% Tween 20 in PBS for 1 hours. Samples were incubated with primary antibody diluted in staining solution (2% normal goat or horse serum 0.2% Tween 20 in PBS) overnight at 4 C°. After 2 hours of washing in 0.2% Tween 20 in PBS, tissue sections were incubated with secondary antibody and DAPI diluted in staining solution for 1 hour at room temperature, followed by washing three times with 0.2% Tween 20 in PBS for 5 min each. Terminal deoxynucleotidyl transferase dUTP nick end labeling (TUNEL) assay was performed using the In Situ Cell Death Detection Kit (Roche) according to manufacturer’s instructions. In brief, following deparaffinization and slide rehydration, slides were treated with a 20 mg/ml Proteinase K (Denville Scientific) solution in PBS for 15 min at room temperature, washed in 0.2% Tween 20 PBS and incubated in TUNEL reaction mixture for 1 hour at 37 C°, then washed prior to mounting. As a positive control for the reaction, we included a slide treated with DNase I for 15 min at room temperature prior to TUNEL reaction mixture incubation. For Hematoxylin and eosin (H&E) staining, sections were fixed in 4% paraformaldehyde for 10 min, washed in PBS and stained in hematoxylin for 8 min and eosin for 1 min. The following primary antibodies were used: Anti-Ki67 (1:100, 14-5698, eBioscience), anti-Keratin 14 (1:1000, 905301, BioLegend), anti-Keratin 10 (1:500, 905401, BioLegend), anti-BrdU (1:100, B-35128, Invitrogen), anti-Lamin A/C (1:100, ab133256, abcam), anti-Ecadherin (1:200, ab16505, abcam), anti-Vinculin (1:50, ab155120, abcam), anti-F-actin (1:100, MA1-80729, Invitrogen), anti-CD3 (1:200, ab5690, abcam), Wheat Germ Agglutinin (WGA) (1:1000, 29022-1, Biotium), and Non-muscle Myosin II A (1:100, 909802, BioLegend). The following secondary antibodies were used: Alexa Fluor 488 Goat anti-Mouse (1:500, A-11001, Invitrogen), Alexa Fluor 594 Goat anti-Rabbit (1:500, A-11012, Invitrogen), Alexa Fluor 488 Goat anti-chicken (1:500, A-110039, Invitrogen), Alexa Fluor 594 Donkey anti-goat (1:500, A-110058, Invitrogen) and Alexa Fluor 647 Donkey anti-goat (1:500, A-21447, Invitrogen). RNAscope experiment was performed by using the RNAscope Multiplex Fluorescent Reagent Kit, version 2 (#323100, Advanced Cell Diagnostic, Newark, CA) with Murine Piezo1 and K14 specific probe (Advanced Cell Diagnostics). All images were acquired using an Olympus BX51 equipped a Hamamatsu Orca CCD camera or a Leica DM6 B equipped with a Leica DFC9000 GT fluorescent camera and a Leica DMC2900 brightfield camera.

### Cell preparation for flow cytometry and sorting

The dorsal skin was dissected, and subcutaneous fat was removed from the skin. To separate epidermal and dermal layers, skin was cut into small pieces and floated, epidermis side up, on 0.25% Trypsin (GIBCO) in PBS for 1 hour at 37 C°. Following the incubation, the epidermis was carefully scraped off and transferred to 10% FBS (fetal bovine serum) in DMEM (GIBCO. Single cell suspensions were obtained by pipetting gently and filtering through 70 mm cell strainers (VWR). Each sample was centriguged and resuspended in 2% FBS in PBS. For cell staining, antibodies were directly added to the cell suspension and incubated for 10 min on ice. Afterward, cells were washed using 2% FBS PBS and resuspended for sorting in 5 mM EDTA PBS. The following antibodies were used for cell sorting experiments: anti-CD49f-APC (1:300, 313615, BioLegend), anti-CD49f-PercP/Cy5.5 (1:300, 313618, BioLegend), anti-CD34-Alexa Fluor 647 (1:50, 560230, BD PharMingen), anti-Sca1-Ly-6A/E-Violet 605 (1:1000, 108133, BioLegend), and DAPI (1:1000, Biotium). Epidermal cells for RNA-sequencing experiments were membrane GFP+, CD49f+, Sca1+, CD34- and DAPI-. Analysis of cell cycle: cells were fixed and permeabilized with BD Cytofix / Cytoperm buffer (BD Pharmingen) on ice after immunofluorescent staining of antibodies specific for cell surface markers; Afterward, resuspended cells were washed using 1X BD Perm/Wash buffer (BD Pharmingen), then cells were stained with intracellular antigen-specific antibodies, including Ki67 antibody (1:100, 14-5698, eBioscience); cells were incubated with DNase-free RNase A to digest doble stranded RNA; At last, to label the toal cellular DNA content for cell cycle analysis, cells were stained with propidium iodide (PI) (1:1000, P3566, Invitrogen). Stained cells were acquired on a flow cytometer. The following antibodies were used for cell cycle analysis: anti-CD49f-FITC (1:200, 313606, BioLegend), anti-Sca1-Ly-6A/E-Violet 605 (1:1000, 108133, BioLegend), anti-Ki67 (1:100, 14-5698, eBioscience), propidium iodide (PI) (1:1000, P3566, Invitrogen), Zombie Kit (1:1000, 423105, BioLegend), and DAPI (1:1000, Biotium). We used Flow cytometry was performed on a BD LSRII cytometer (BD Biosciences), sorting on a BD FACS Aria II sorter (BD Biosciences). Flow cytometry data was collected and exported using BD FACs Diva software (BD Biosciences) and analyzed and plotted using FlowJo software.

### RNA sequencing/data analysis

p63CreER; Piezo1fl/fl; R26-mTmG (Piezo1 cKO) mice were administered tamoxifen injections (3 times, 20ug/dose) at P45-P52 to activate cre recombination and delete Piezo1 gene expression in basal layer keratinocytes. Basal epidermal cells were isolated from Piezo1 cKO and control mouse back skin using flow sorting. RNA was extracted using the RNeasy Mini Kit (QIAGEN) following manufacturer’s instructions. The libraries for sequencing were prepared using the method described before90. Briefly, RNA-seq libraries were prepared at the same time for all samples belonging to a single experimental cohort to reduce batch effects. All RNA-seq libraries were prepared using the NEBNext Poly(A) mRNA magnetic isolation module, followed by NEBNext Ultra Directional RNA library preparation kit for Illumina. Library quality was checked by Agilent BioAnalyzer 2100, and libraries were quantified using the Library Quant Kit for Illumina. Libraries were then sequenced using a NextSeq500 platform [75–base pair (bp) single-end reads]. All RNA-seq was aligned using RNA STAR 91 under default settings to Mus musculus GRCm38 fragments per kilobase per million mapped fragments, and differential expression analysis were performed using DESeq292. Statistical significance was obtained using an adjusted P value generated by DESeq2 of less than 0.05. Replicates were generated from 3 control and 3 Piezo1 cKO mice, isolated and sorted at different time. All GO analyses were performed using PANTHER at http://pantherdb.org/ to determine statistically overrepresented GO terms using Fisher’s exact test under the “biological process” category. P values for GO terms are false discovery rate statistics. The top 12 plotted GO terms represent the GO terms with the highest fold enrichment under PANTHER’s default hierarchical clustering categorization. GO term figures are generated using ggplot2.

### Engineered human skin xenografts

Organotypic skin grafts containing human primary keratinocytes were performed as previously detailed methods^107–110^. Primary human keratinocytes that provided from SBDRC Core B were initiated from adult human skin and plated directly for experiments. Human primary keratinocytes that genetically encoded with fluorescent reporter genes were cultured in a 1:1 mixture of Keratinocyte Growth Media (KGM) and Keratinocyte Media 50/50 (Gibco) containing 2% FBS,

mM calcium chloride, 100 nM Et-3 (endothelin 3), 10 ng/mL recombinant human stem cell factor, and 4.5 ng/mL recombinant basic fibroblast growth factor. Briefly, primary human keratinocytes were transduced with K14H2B-PAGFP and Lifeactin-EGFP. Transduced human keratinocytes (10 × 10^5^) were suspended in 80 µl mixture media, seeded onto the decellularized human dermis that Ridky lab and SBDRC Core B generously provided, and incubated at 37 C° for 4 days at the air-liquid interface to establish organotypic skin. Organotypic skin tissues were grafted onto 5-to 7-week-old female ICR SCID mice (Taconic) according to an International Animal Care and Use Committee (IACUC) – approved protocol at the University of Pennsylvania. Mice were anesthetized in an isoflurane chamber, and murine skin was removed from the upper dorsal region of the mouse. Organotypic human skin was trimmed to a uniform 11 mm by 11 mm square and grafted onto the back of the mouse with individual interrupted 6-0 nylon sutures. Mice were dressed in Bactroban ointment, Adaptic, Telfa pad, and Coban wrap. Dressings were removed 2 weeks after grafting. Mechanical treatments were performed 2 months after grafting.

### Calcium imaging data analysis

To investigate the cellular response to mechanical forces, calcium dynamics were captured in individual epidermal cells within basal layer of epidermis *in vivo* using *R26-CAG-GCaMP6s*; *R26-mTmG* mice. At both rest and stressed condition, time lapses of one z plane (the basal layer of epidermis) were recorded. Time-lapse images were analyzed and plotted using the following steps. First, raw image stacks were imported into FIJI (ImageJ, NIH); To measure the calcium dynamics in the same cells in response to mechanical forces, region of interest (ROIs) on the cells that responded to stresses were chose and segmented manually based on tomato fluorescent labeled cell membrane. After setting multiple RO1s using ROI manager, the mean fluorescence intensity (GFP) of each segmented cell was measured and normalized to basal level for all segmented cells and frames.

### Digital data analysis

Raw image files from two-photon imaging were imported into FIJI (Image J NIH) or to Imaris (Bitplane) for analysis. Individual optical planes and sequential optical sections were used to assemble figures as previously described. Fig. 1D and Extended Data Fig. 2A were created using Imaris software. To quantify population clonal dynamics and high-resolution optical sections were obtained sequentially and used to construct 3-dimensional tiled views of the epidermis. The same mice were then re-imaged using identical acquisition parameters. For single-cell lineage tracing, individual high-magnification serial optical sections were obtained for each traced clone and 3-dimensional analysis was performed once the entire imaging time course was competed to analyze the state of basal and suprabasal cells in each time point. Departure of a cell from the basal layer and subsequent upward transit was scored as differentiation while continuous increase in the basal cell number was scored as self-renewal. Clone measurements were performed manually. Images shown in figures typically represent maximum projections or single optical sections selected from the z stacks unless otherwise specified.

### Electron microscopy

Trimmed skin samples were fixed in 2% glutaraldehyde and 2% paraformaldehyde in 0.1M sodium cacodylate buffer pH7.4 overnight. Samples were submitted to and processed by Electronic Microscopy Resource Lab (EMRL), a core facility at UPenn. Images were taken using Morada CCD and iTEM (Olympus) software.

### Statistics and reproducibility

Data are expressed as column bar graphs and scatter plots. An unpaired Student’s t-test was used to analyze data sets with two groups and *P < 0.05 to ****P < 0.0001 indicated a significant difference. Statistical calculations were performed using the Prism software package (GraphPad). No statistical method was used to predetermine sample size.

---

**Supplementary Figure 1.**
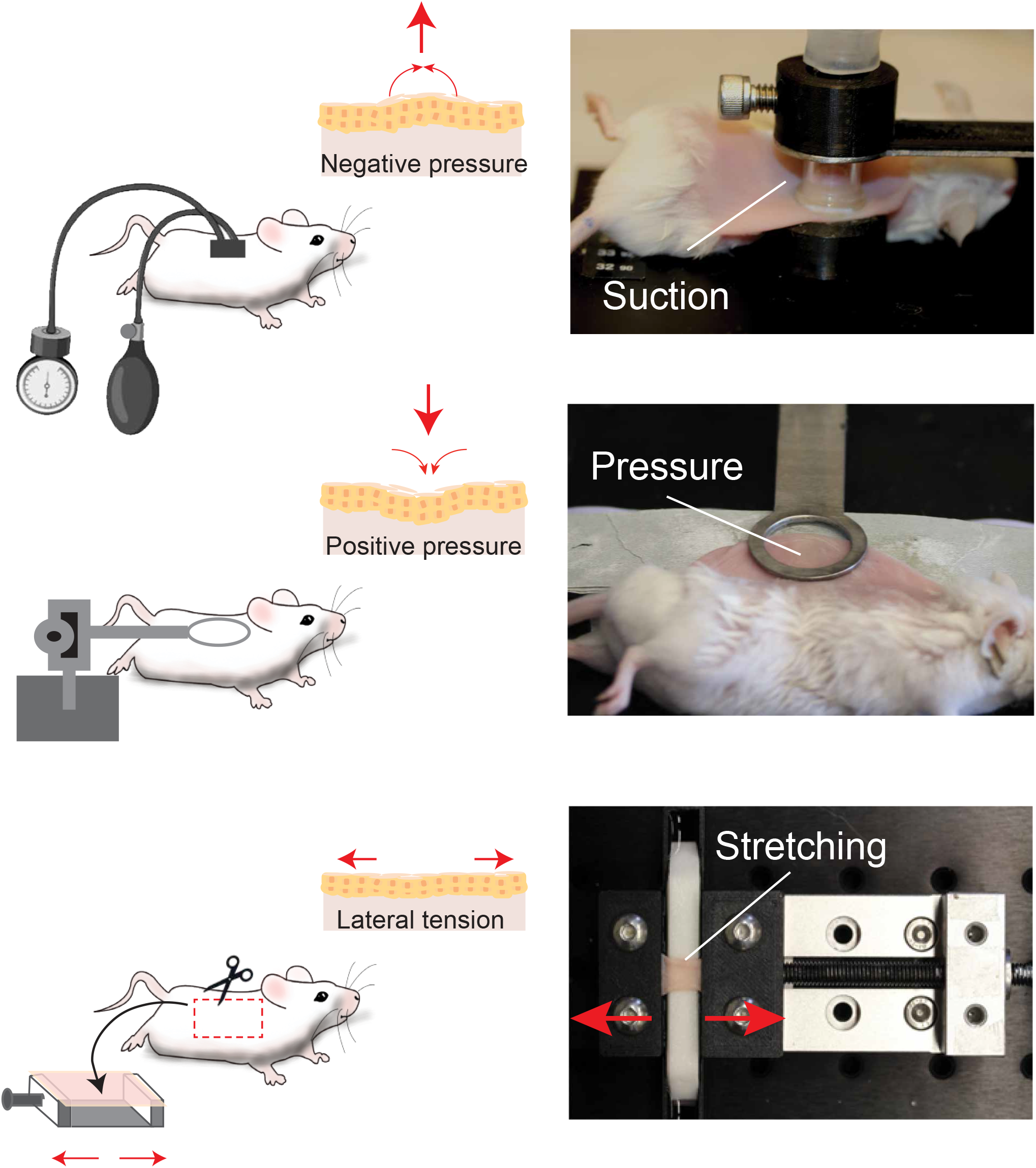
Schematic representations of the *in vivo* and *ex vivo* assays employed in this study to analyze the effect of acute mechanical forces applied on the mouse skin, by intravital imaging.

**Supplementary Figure 2.**
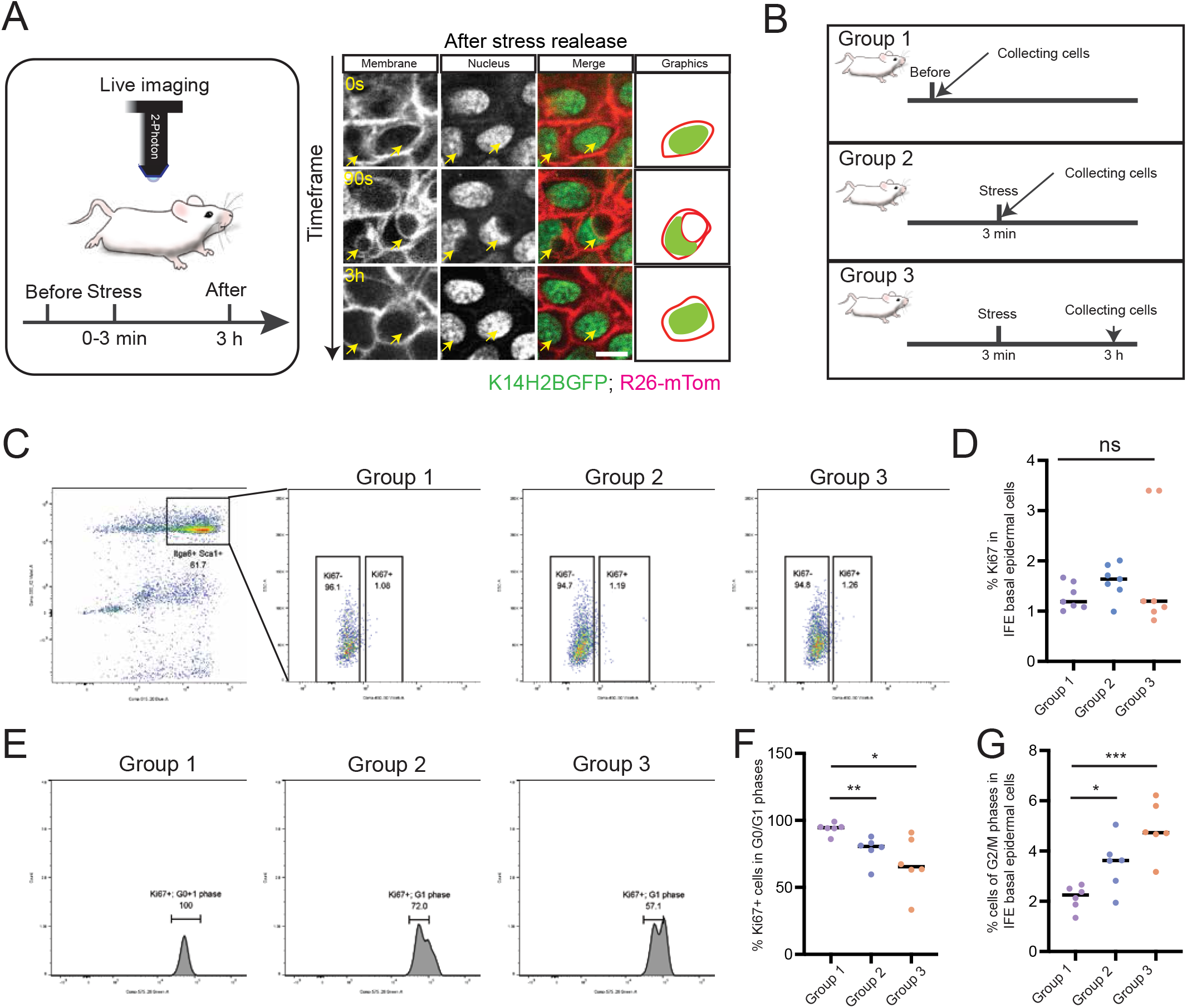
(A) Representative time frames of basal epidermal cells, visualizing the stress vesicle resorption after force removal. Yellow arrows indicate the locations of stress vesicles. Scale bars, 5 µm. (B) Schematic of the experimental design to isolate epidermal cells from mouse skin before and after mechanical stress. (C) Representative Flow cytometry data showing the percentage of Ki67+ cell in the interfollicular epidermis (Itga6+ and Sca1+) that were collected from normal skin, stressed skin, and the recovered skin (3 hours after treatment). (D) Quantification of Ki67 rates in distinct samples. n= 6 mice for each group, p=0.0.0739 (ns) and 0.3447 (ns). (E) Representative data of cell phase shifts in Ki67+ cells, showing G1/S to G2/M transition. (F) Quantification of the rates of Ki67+ cells that still stayed in G1 phase. n= 6 mice for each group, p=0.0048 (**) and 0.0109 (*). (G) Quantification of the rates of Ki67+ cells that stayed in G2/M phase. n= 6 mice for each group, p=0.0.0172 (*) and 0.0002 (***).

**Supplementary Figure 3.**
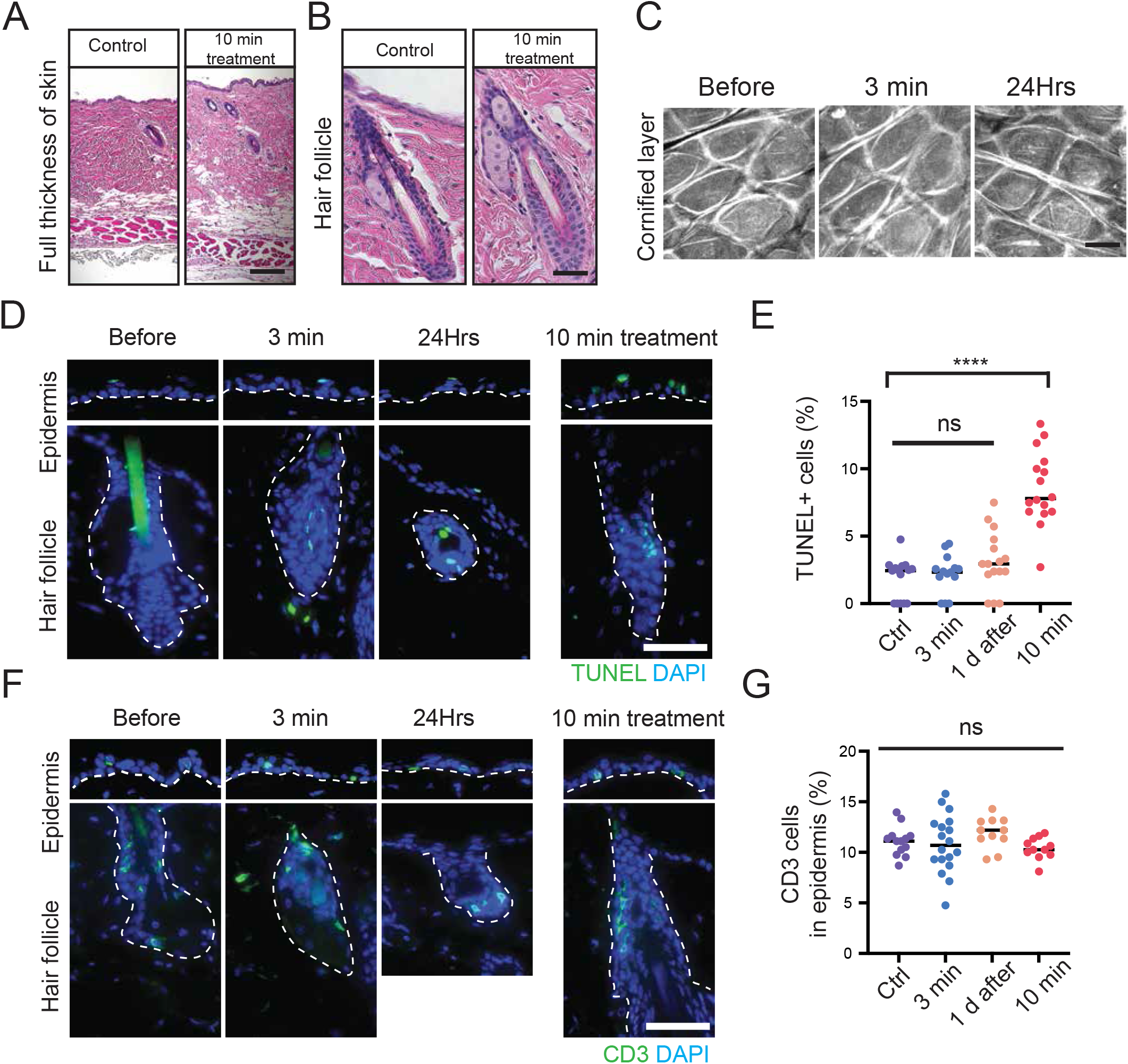
(A) H&E staining of full thickness back skin samples that were collected from control and stressed mice. Scale bars, 500 µm. (B) H&E staining of normal and stressed skin samples with magnified hair follicle area. Scale bars, 50 µm. (C) Representative live images at the plane of the cornified layer of the epidermis captured at the indicated time points. Scale bars, 10 µm. (D) Representative images of TUNEL staining. A white dashed line demarcates the border between the epidermis and dermis. Scale bars, 20 µm. (E) Quantification of the rates of apoptotic cells in samples collected at the indicated time points. n = 15 areas from 3 mice (normal), 15 areas from 3 mice (3 min treatment), 16 areas from 3 mice (1 day after treatment), and 16 areas from 3 mice (10 min treatment), p= 0.7290 (ns), 0.0725 (ns) and p < 0.0001 (****). (F) Immunofluorescence staining of T-cells (CD3) in normal, stressed (3 min treatment) and recovered epidermis (1 day after treatment), and highly stressed epidermis (10 min treatment). Scale bars, 20 µm. (G) Quantification of CD3+ immune cells in distinct samples that were collected at the indicated time points. n = 13 areas from 3 mice, 18 areas from 3 mice, 11 areas from 3 mice, and 11 areas from 3 mice, p= 0.7759 (ns), 0.1841(ns), and 0.1956 (ns).

**Supplementary Figure 4.**
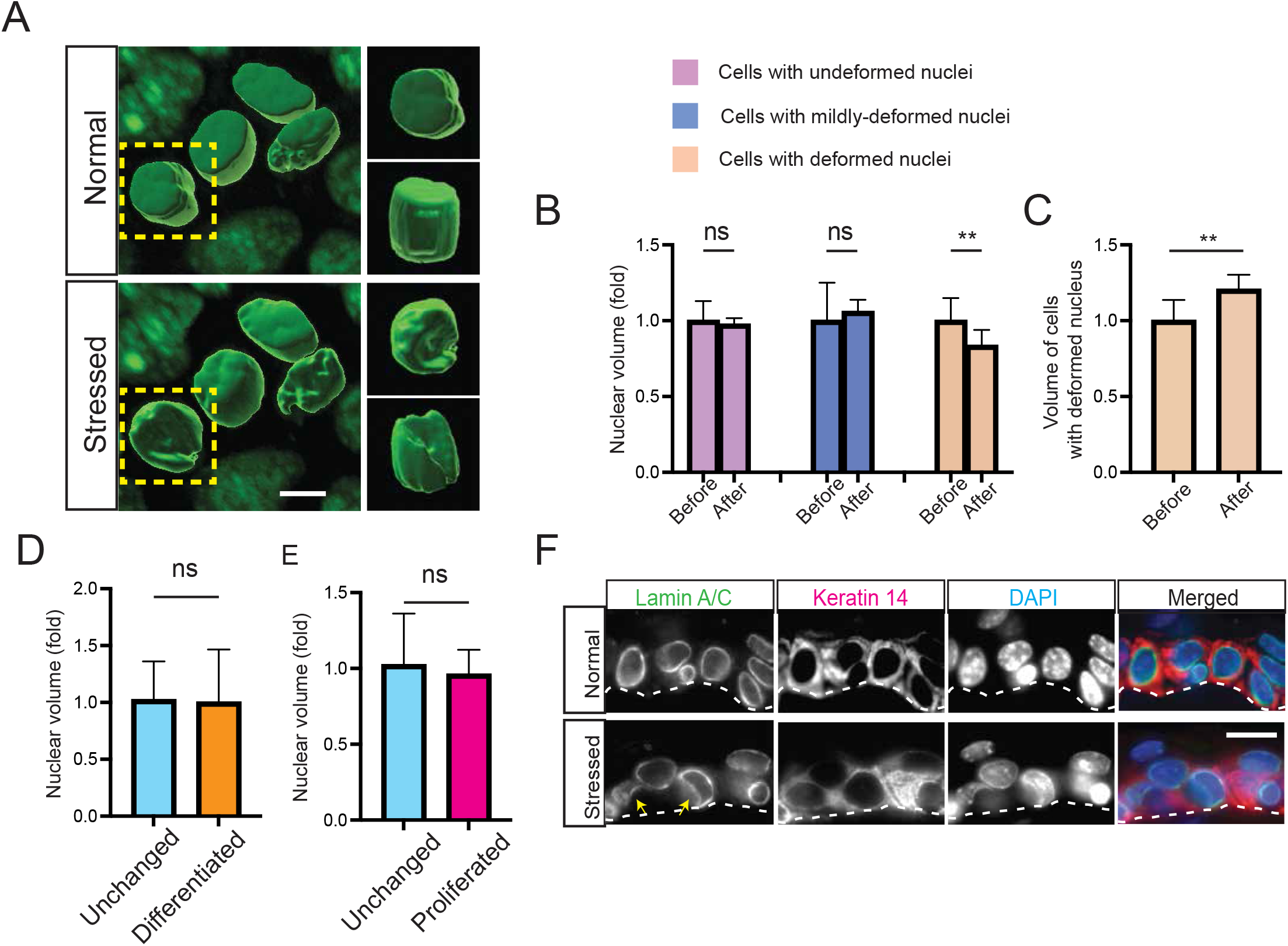
(A) High-magnification 3-dimensional surface rendering demonstrating the nuclear deformation under stressed condition. Scale bars, 2 µm. (B) Quantification of nuclear volume before and after mechanical force application. n = 10, 16, and 14 cells from 3 mice, p = 0.7025 (ns), 0.3713 (ns), and 0.0023 (**). (C) Quantification of cell volume. n = 28 cells from 3 mice, p = 0.0072 (**). (D) and (E) Quantification of nuclear volume in differentiated (left graph) or proliferated (right graph) cells compared to undifferentiated cells. n= 18 undifferentiated cells, 15 differentiated cells, and 12 proliferated cells from 3 mice, p = 0.9058 (ns) and 0.7597 (ns). (F) Co-localization by immunofluorescence of Lamin A/C (green) and Keratin 14 (red) in normal and stressed epidermis. Scale bars, 5 µm.

**Supplementary Figure 5.**
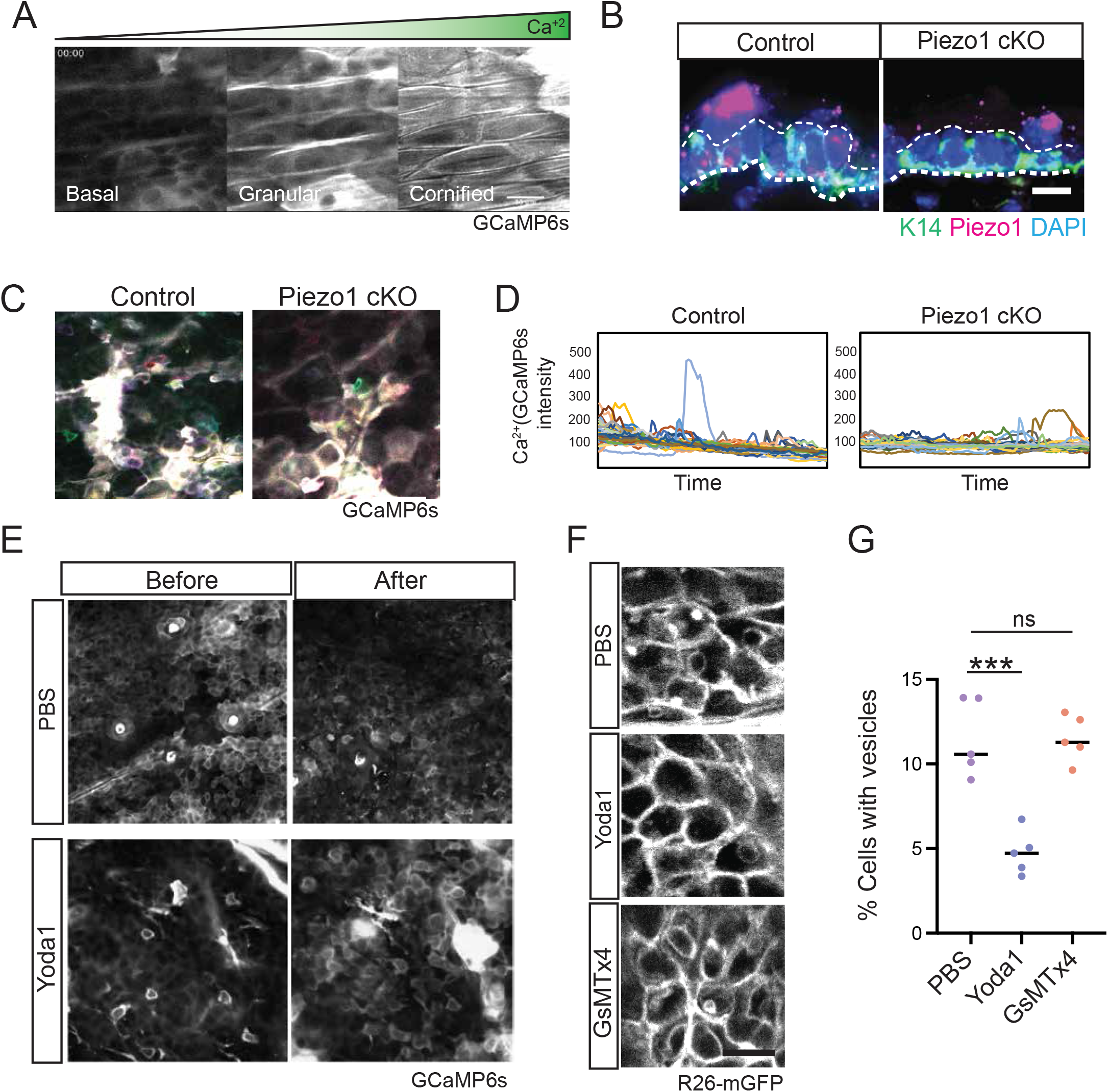
(A) Representative planes from serial optical sectioning of the live mouse epidermis showing the calcium gradient from the basal to suprabasal layers. (B) Representative RNA-scope staining indicating the efficiency of Piezo1 gene deletion epidermal cells. Scale bars, 10 µm. (C) Maximum intensity projection of calcium signals from timelapse imaging of control and Piezo1 cKO mouse skin under normal conditions. Individual frames were pseudo-colored and projected into a single image to illustrate areas of persistent calcium signal (white). Scale bars, 50 µm. (D) Quantification of calcium signals from (C). (E) Representative images of *in vivo* imaging of calcium flux after Yoda1 (Piezo1 activator) treatment. (F) Representative images of stress vesicle formation after treatment with PBS, Yoda1(Piezo1 agonist), or GsMTx4 (Piezo1 inhibitor). Scale bars, 20 µm. (G) Quantification of the number of cells with vesicle formation after the treatment of PBS, Yoda1, and GsMTx4. n= 5 mice for each group, p = 0.004 (***) and 0.9940 (ns).

**Supplementary Figure 6.**
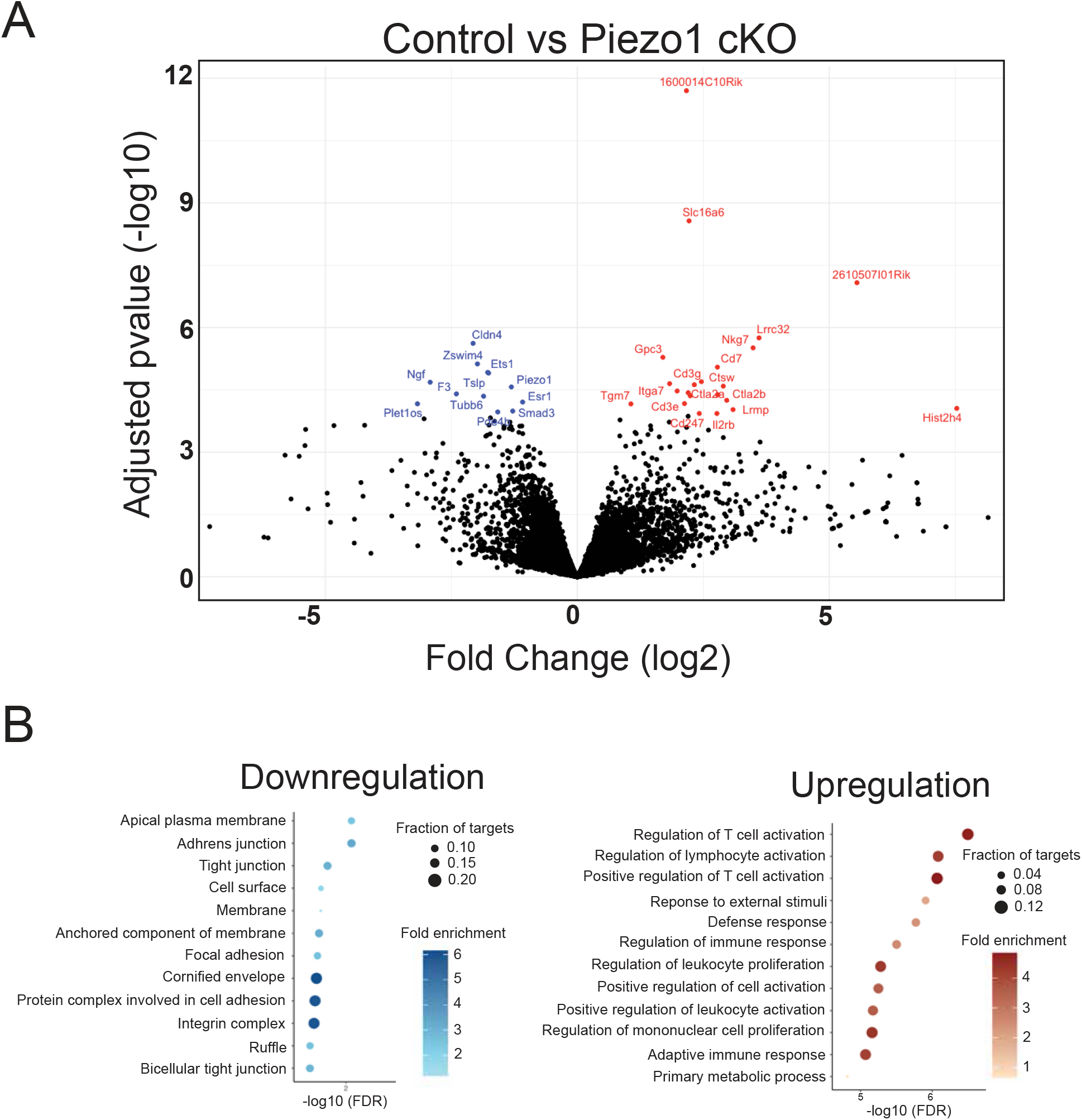
(A) Volcano plot of RNA sequencing data, comparing the transcripts in the basal epidermal cells that are collected from control and Piezo1 cKO. Red dots represent upregulated genes, and blue dots represent downregulated genes; adjusted p < 0.05. (B) Gene Ontology analysis of genes upregulated genes and downregulated genes from the RNA-seq analysis.

**Supplementary Figure 7.**
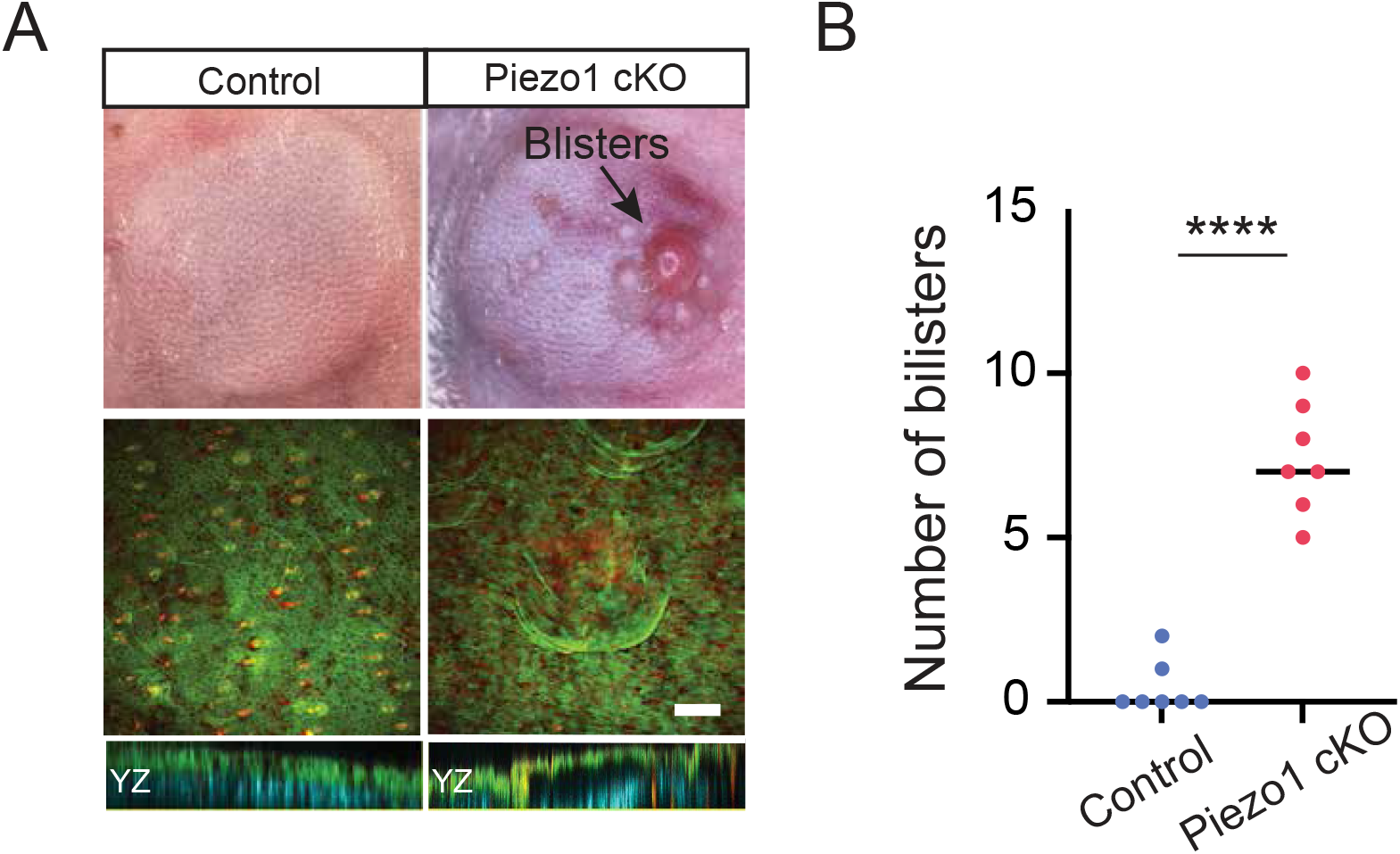
(A) Examples of mouse back skin imaged by brightfield (top panel) and two-photon (lower panel) microscopy after 10 min of applied negative pressure. Scale bars, 100 µm. (B) Quantification of the number of blisters on control and Piezo1 cKO back skin after 10 min of applied negative pressure. n= 7 pairs of mice, p < 0.0001.

## Notes

### Competing Interest Statement

The authors have declared no competing interest.

### Summary of Updates

Correct minor typographical errors

